# REN-former prioritizes candidate regulators of kidney disease-state transitions through single-cell foundation modeling and human genetics

**DOI:** 10.64898/2026.09.10.746481

**Authors:** Imari Mimura, Sosuke Hosokawa, Yosuke Hirakawa, Toshinaru Kawakami, Yu Kurata, Masamichi Ito, Tetsuhiro Tanaka, Satoshi Kodera, Norihiko Takeda, Masaomi Nangaku

**Affiliations:** Division of Nephrology and Endocrinology, Graduate School of Medicine, The University of Tokyo, Tokyo, JAPAN; Organ Pathophysiology Program, The University of Tokyo, Tokyo, JAPAN; Department of Information and Communication Engineering, Graduate School of Information Science and Technology, The University of Tokyo, Tokyo, JAPAN; Department of Cardiovascular Medicine, Graduate School of Medicine, The University of Tokyo, Tokyo, JAPAN; Department of Nephrology, Rheumatology and Endocrinology, Tohoku University Graduate School of Medicine

**Keywords:** Geneformer, single-cell RNA sequencing, foundation model, proximal tubule, in silico perturbation, AKI-to-CKD transition

## Abstract

**Background:** Acute kidney injury (AKI)-to-chronic kidney disease (CKD) transition is associated with dynamic changes in tubular cell state. However, conventional single-cell transcriptomic analyses primarily identify genes differentially expressed between disease states and do not directly evaluate genes associated with directional state transitions.

**Methods:** We developed REN-former by fine-tuning Geneformer using the GSE183276 single-cell RNA-sequencing dataset from 45 participants representing Normal Reference, AKI, and CKD. In silico gene deletion and overexpression analyses estimated directional transcriptomic shifts in proximal tubular cells. Candidates were evaluated using summary-data-based Mendelian randomization (SMR), colocalization and expression analysis in additional KPMP participants not included in GSE183276.

**Results:** REN-former achieved recall values of 0.99, 0.80, and 0.79 for Normal Reference, AKI, and CKD, respectively. In silico perturbation analyses identified distinct gene programs associated with transitions from Normal Reference to AKI, from Normal Reference to CKD, from AKI to CKD, and from CKD to Normal Reference. Conventional analysis showed metabolic suppression and increased inflammatory and stress-response activation. Six genes—*IFITM3, CALR, TTR, CALM1, MUC13,* and *RPL13*—met the prespecified SMR, HEIDI, and directional-concordance criteria, and colocalization supported *IFITM3, CALR, TTR,* and *CALM1*.In additional KPMP data, *IFITM3* was higher, whereas *TTR* and *CALM1* were lower, in CKD proximal tubules; *CALR* did not differ significantly. The observed expression changes were concordant with the REN-former-predicted directions for *TTR* and *CALM1* but discordant for *IFITM3*.

**Conclusion:** REN-former provides a framework for prioritizing candidate regulators of kidney disease-associated cell states by integrating predicted perturbation effects with human genetic and transcriptomic evidence.

## Introduction

Chronic kidney disease (CKD) is a major global health problem associated with substantial morbidity, mortality, and healthcare burden. In 2017, approximately 700 million people worldwide were estimated to have CKD, and approximately 1.2 million deaths were directly attributable to CKD^1^. Acute kidney injury (AKI) is also a common clinical syndrome and is increasingly recognized not only as an acute event but also as an important risk factor for subsequent CKD and kidney failure^2–6^. Although kidney function may recover after AKI, some patients experience persistent kidney dysfunction and progressive loss of kidney function^7^. The molecular mechanisms that distinguish successful repair from progression toward CKD remain incompletely understood. Identifying the genes and biological programs that regulate the transition from AKI to CKD is therefore important for developing strategies to promote adaptive repair and prevent chronic kidney disease progression^6, 8–10^.

Proximal tubular cells are particularly susceptible to ischemic, toxic, and inflammatory injury because of their high metabolic demands and dependence on mitochondrial energy production. Following AKI, surviving differentiated tubular epithelial cells can proliferate and contribute to reconstruction of the damaged tubular epithelium^11^. However, after severe or persistent injury, a subset of proximal tubular cells may fail to return to a differentiated state and instead acquire proinflammatory and profibrotic transcriptional programs. Single-cell transcriptomic studies have identified these failed-repair proximal tubular cells and implicated their persistence in maladaptive repair and the transition from AKI to CKD^12–14^. Subsequent single-nucleus studies have further demonstrated that proximal tubular cells occupy multiple dynamic states during injury and repair rather than forming uniform healthy and injured populations^15, 16^. Comprehensive human kidney atlases have similarly revealed extensive heterogeneity among healthy and disease-associated epithelial states and have provided a framework for investigating cellular programs underlying kidney injury, repair, and chronic disease^17^.

Conventional single-cell RNA-sequencing analyses typically identify cell populations, disease-associated cell states, differentially expressed genes, and enriched biological pathways. These approaches are highly effective for characterizing transcriptional differences observed between disease states. However, they primarily describe static expression patterns and do not directly evaluate how perturbation of an individual gene may shift a cell from one transcriptomic state toward another. Recently, foundation models pretrained on large collections of single-cell transcriptomes have provided an opportunity to learn context-dependent relationships among genes and transfer this information to specific biological tasks. Several computational models, including scGen, scGPT, and scFoundation, have been developed to learn patterns from large single-cell RNA-sequencing datasets and to predict cellular responses under different conditions^18–20^. Among these models, Geneformer provides an in silico perturbation framework that can be used to estimate how changes in individual genes may affect cellular states.

Geneformer is a Transformer-based foundation model that represents each cell using a rank-value encoding of expressed genes. In this study, we used the expanded Geneformer V2 model, which was pretrained on approximately 95 million human single-cell transcriptomes.^21^ It represents each cell as a sequence of expressed genes ordered by their normalized expression ranks, rather than as a conventional gene-by-cell expression matrix, enabling the model to learn context-dependent relationships among genes. In this rank-based representation, gene overexpression was modeled by moving the perturbed gene to the highest rank in the cell’s gene sequence. This representation supports transfer learning, gene-network analysis, and in silico perturbation in limited-data settings^22^. However, its application to directional transitions among Normal Reference, AKI, and CKD tubular-cell states remains largely unexplored, although we previously applied a related approach to pulmonary arterial hypertension.^23^ Furthermore, it is unknown whether genes prioritized through foundation-model-based perturbation analysis are supported by conventional transcriptomic analyses and human genetic associations with kidney function.

In this study, we fine-tuned Geneformer with a publicly available human kidney single-cell RNA-sequencing dataset containing Normal Reference, AKI, and CKD samples. We called the resulting kidney-specialized model REN-former. We used in silico gene deletion and overexpression in proximal tubular cells to identify genes associated with the Normal Reference-to-AKI, AKI-to-CKD, Normal Reference-to-CKD, and CKD-to-Normal Reference transitions. We also performed conventional gene expression and pathway analyses using proximal tubular cells from the same dataset. Finally, we combined REN-former candidates with kidney eQTL and eGFR GWAS data using SMR and colocalization analyses. Together, these approaches were used to identify genes and biological programs associated with proximal tubular-cell state changes in AKI and CKD.

## Methods

### Single-cell RNA-sequencing dataset and preprocessing

We analyzed the processed GSE183276^17^ human kidney scRNA-seq dataset, comprising 109,741 cells and 37,080 genes from 18 Normal Reference, 12 AKI, and 15 CKD participants. The Normal Reference group comprised living-donor kidney biopsy samples from 18 participants with eGFR >60 mL/min/1.73 m² and no history of diabetes or hypertension. The AKI group comprised KPMP kidney biopsy samples from participants with KDIGO stage 1 (*n* = 1), stage 2 (*n* = 5), or stage 3 (*n* = 6) AKI. Histopathological findings were heterogeneous; acute tubular injury was present in 10 of the 12 biopsies, including one with acute interstitial nephritis. Diabetic kidney disease and hypertensive CKD samples were combined into the CKD group. Original cell-type annotations and UMAP coordinates were retained, and cells annotated as proximal tubules were used for proximal-tubule analyses. Additional details are provided in the Supplementary Methods.

### Gene identifier mapping and tokenization

The processed data were converted into AnnData format. Gene symbols were converted to Ensembl gene identifiers using GENCODE release 43, and unmapped genes were excluded. For each cell, genes were ranked by expression and converted into Geneformer-compatible tokens, while participant, disease-state, and kidney-structure information was retained.^22^

### REN-former construction and participant-level data split

We used the pretrained Geneformer V2 model GF-12L-95M-i4096,^21^ a 12-layer model pretrained on approximately 95 million human single-cell transcriptomes with a maximum input length of 4,096 genes. We created REN-former by fine-tuning this model to classify individual kidney cells as Normal Reference, AKI, or CKD using the Geneformer cell-classification framework. REN-former was fine-tuned for one epoch with a learning rate of 2 × 10⁻⁵, a batch size of 16, and a random seed of 42. The first four Transformer layers were kept fixed during training. The data were divided by participant into mutually exclusive training, validation, and test sets containing 34, 5, and 6 participants, respectively. All cells from each participant were retained within a single partition. The disease-group composition of each set is provided in the Supplementary Methods. The test set was used for final model evaluation and in silico perturbation analyses.

### Model evaluation

We evaluated the saved REN-former model using the labeled test set, which had not been used for model training. We used Geneformer’s evaluate_saved_model function with a forward batch size of 100 and 16 parallel processes. Model performance was shown using a confusion matrix and the predicted probabilities for each disease group. For each disease group, recall was calculated as the proportion of cells correctly assigned to that group. For example, AKI recall was the proportion of true AKI cells predicted as AKI. We also plotted the predicted probabilities for all three groups according to the true label of each cell.

### In silico deletion and overexpression analyses

We next used REN-former to test the predicted effect of changing one gene at a time. These in silico perturbation analyses were performed with Geneformer’s InSilicoPerturber framework using the test set. Each analysis had a starting state and a predefined goal state.

Deletion analyses examined the Normal Reference-to-AKI, AKI-to-CKD, and Normal Reference-to-CKD transitions. Overexpression analyses examined the Normal Reference-to-CKD and AKI-to-CKD transitions. We also performed reverse analyses to identify changes that moved CKD-like cells toward a Normal Reference-like state, using both deletion and overexpression.

Each gene was tested separately. Deletion was simulated by removing the gene, whereas overexpression was simulated by moving the gene toward the top of the ranked gene list. For each gene, REN-former assessed whether the simulated perturbation moved cells closer to the target disease state. The resulting Shift_to_goal_end value was used to rank genes. Each analysis included up to 300 cells from the starting state. Deletion analyses were performed in both all eligible kidney cells and proximal tubular cells for the Normal Reference-to-AKI, AKI-to-CKD, and Normal Reference-to-CKD transitions. Biological interpretation focused mainly on proximal tubular cells.

### Perturbation statistics and candidate gene selection

Statistical testing of the perturbation results was performed with Geneformer’s InSilicoPerturberStats using mode=“goal_state_shift”. For each gene, the output included the direction and size of the shift toward the goal state, a comparison with random perturbations, a nominal P value, an FDR-adjusted P value, and the number of cells in which the gene was tested. A gene was considered a significant candidate when its perturbation produced a positive shift toward the goal state and its FDR-adjusted P value was below 0.05. The number of tested cells differed among genes because a gene could be perturbed only in cells in which that gene token was present.

For deletion analysis, we compared the significant genes found in all kidney cells with those found in proximal tubular cells. This comparison was performed separately for the Normal Reference-to-AKI, AKI-to-CKD, and Normal Reference-to-CKD transitions. Shared and analysis-specific genes were shown using Venn diagrams.

### Participant-level differential expression and Hallmark pathway analysis

Proximal-tubule UMI counts from GSE183276 were aggregated by participant to generate pseudobulk profiles, and participants with fewer than 20 proximal-tubule cells were excluded. The analysis included 24,985 cells from 18 Normal Reference, 12 AKI, and 15 CKD participants. Differential expression was analyzed using robust quasi-likelihood models in edgeR, with disease group and, where possible, sex included in the model.^24, 25^ Genes with FDR <0.05 and absolute log2 fold change ≥0.5 were considered differentially expressed. Preranked Hallmark gene set enrichment analysis was performed using fgsea and MSigDB Hallmark gene sets, with Benjamini–Hochberg-adjusted *P* <0.05 considered significant.^26, 27^ Additional details are provided in the Supplementary Methods.

### Gene Ontology enrichment analysis

GO Biological Process enrichment analysis was performed separately for each transition, perturbation type, and cell population.^28^ Directional candidates were defined by a positive shift toward the goal state and an FDR below 0.05. All genes statistically tested in the corresponding perturbation analysis were used as the background.

Gene symbols were mapped to Entrez Gene identifiers, and enrichment was tested using clusterProfiler.^29^ GO terms with a Benjamini–Hochberg-adjusted *P* value below 0.05 and at least three candidate genes were considered significant. Redundant terms were reduced using Wang semantic similarity with a cutoff of 0.7.^30^ Additional details are provided in the Supplementary Methods.

### Summary-data-based Mendelian randomization (SMR) and colocalization analysis

We performed summary-data-based Mendelian randomization (SMR).^31^ SMR combines expression quantitative trait locus (eQTL) data with genome-wide association study (GWAS) data to identify genes whose genetically predicted expression is associated with a trait. We used tubulointerstitial eQTL summary statistics from NephQTL.^32^ These data were combined with summary statistics from a large multi-ancestry eGFR GWAS.^33^ Variants with a minor allele frequency below 0.05 were excluded, and European reference data were used to estimate linkage disequilibrium.

Genes were taken forward for colocalization when they met the Bonferroni-corrected SMR threshold (*P* < 0.05/199), showed no significant heterogeneity by the HEIDI test (HEIDI *P*> 0.05), and had an SMR effect direction concordant with the corresponding REN-former-predicted perturbation direction. Concordance was defined as an SMR effect indicating higher eGFR in the expression direction predicted to shift cells toward a Normal Reference-like state, or lower eGFR in the direction predicted to shift cells toward a CKD-like state. For example, when gene deletion was predicted to shift cells toward CKD, the corresponding SMR effect was required to indicate higher eGFR with increased genetically predicted expression. Colocalization analysis was then performed using the coloc R package.^34^ Variants within ±250 kb of each gene’s transcription start site were analyzed to assess whether the tubulointerstitial eQTL and eGFR GWAS associations were consistent with a shared causal variant. A posterior probability for hypothesis 4 (PP.H4) ≥ 0.80 was considered strong evidence of colocalization.

### Expression analysis in non-overlapping KPMP participants

Additional KPMP scRNA-seq and snRNA-seq data were analyzed after excluding participants overlapping or potentially overlapping GSE183276 and those with fewer than 20 proximal-tubule cells. Counts were aggregated by participant to generate pseudobulk profiles and analyzed using robust quasi-likelihood models in edgeR. Models included disease and, where available, sex and age; the snRNA-seq analysis additionally adjusted for assay profile. The scRNA-seq analysis was primary, and snRNA-seq was used for replication. Log2 fold changes were defined as CKD relative to Normal Reference. *P* values were adjusted across the four prespecified genes using the Benjamini–Hochberg method, with FDR <0.05 considered significant.

### Data Availability Statement

The GSE183276 dataset is available from the Gene Expression Omnibus. Expanded KPMP scRNA-seq and snRNA-seq data are available from the KPMP data repository subject to a data use agreement. Tubulointerstitial eQTL summary statistics were obtained from NephQTL. Multi-ancestry eGFR GWAS summary statistics reported by Liu et al.^33^ are available from Figshare (DOI: 10.6084/m9.figshare.26299093) under a CC BY 4.0 license. Analysis code, software environment information, and derived results underlying the figures and tables are available from the corresponding author upon reasonable request.

## Results

### REN-former captures transcriptomic differences among Normal Reference, AKI, and CKD kidney cells

To construct a kidney-specialized Geneformer model, we used a publicly available human kidney single-cell RNA-sequencing dataset comprising 45 participants, including 18 Normal Reference participants, 12 patients with acute kidney injury (AKI), and 15 patients with chronic kidney disease (CKD) (Figure 1A and Table 1). The processed dataset contained 109,741 cells, including 21,650 Normal Reference cells, 35,777 AKI cells, and 52,314 CKD cells after diabetic kidney disease and hypertensive CKD samples were combined into a single CKD category. Visualization using the reference UMAP coordinates provided with the original dataset showed that cells from Normal Reference, AKI, and CKD kidneys were distributed across the major kidney cell populations (Figure 1B and 1C). Disease groups were not separated into entirely distinct cell clusters, consistent with the presence of disease-associated transcriptional variation within shared kidney cell populations.

**Figure 1.**
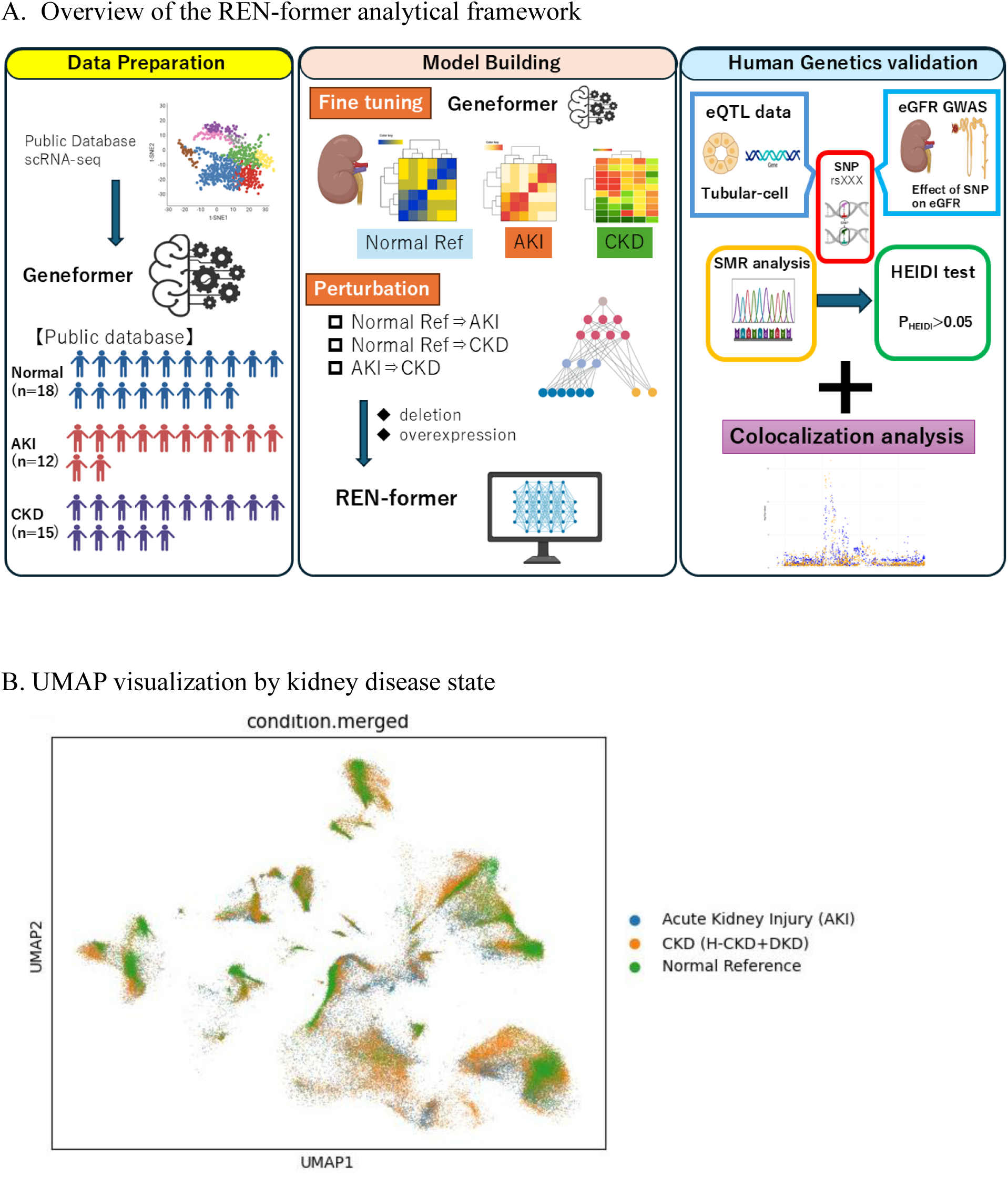
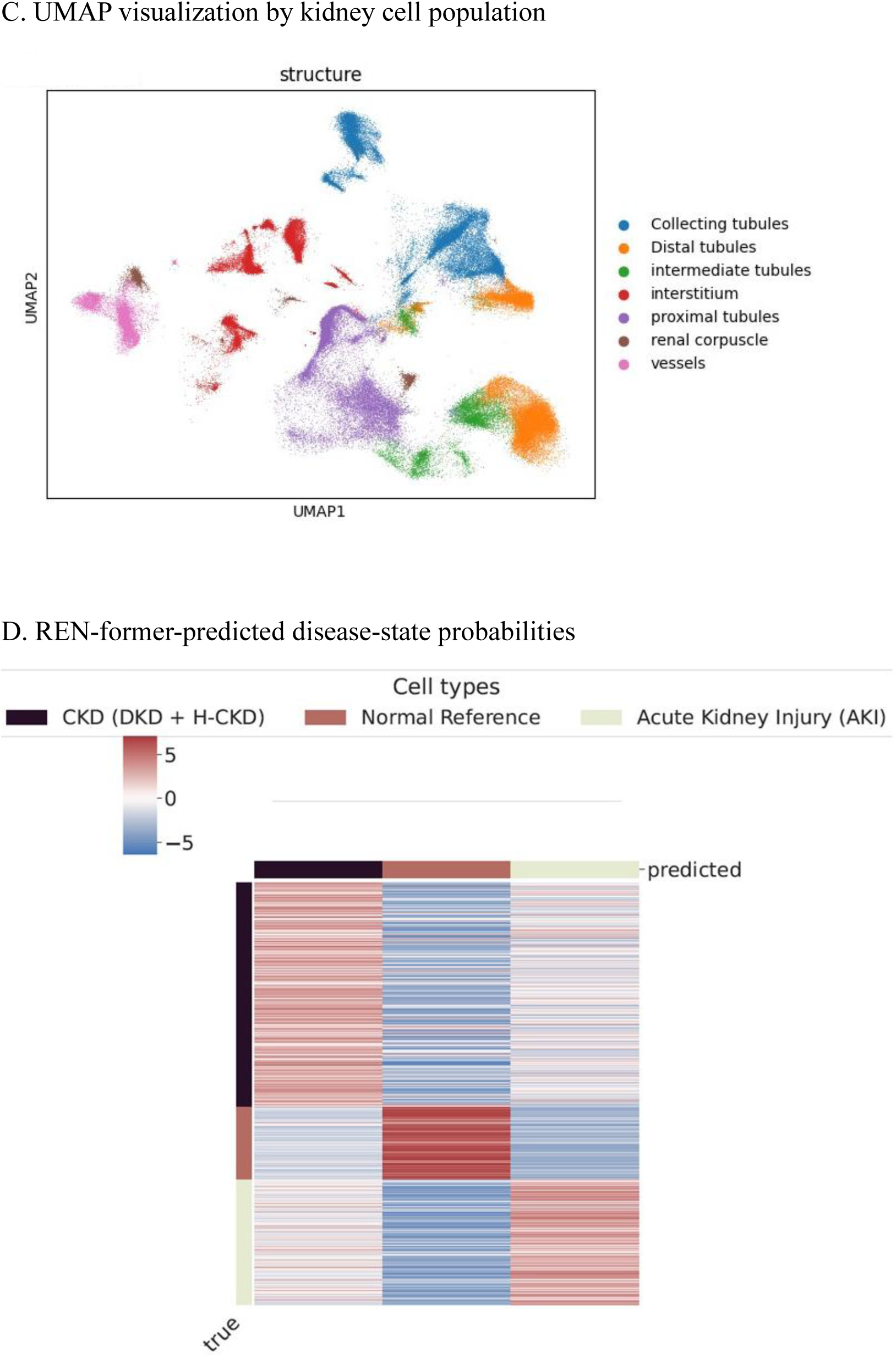
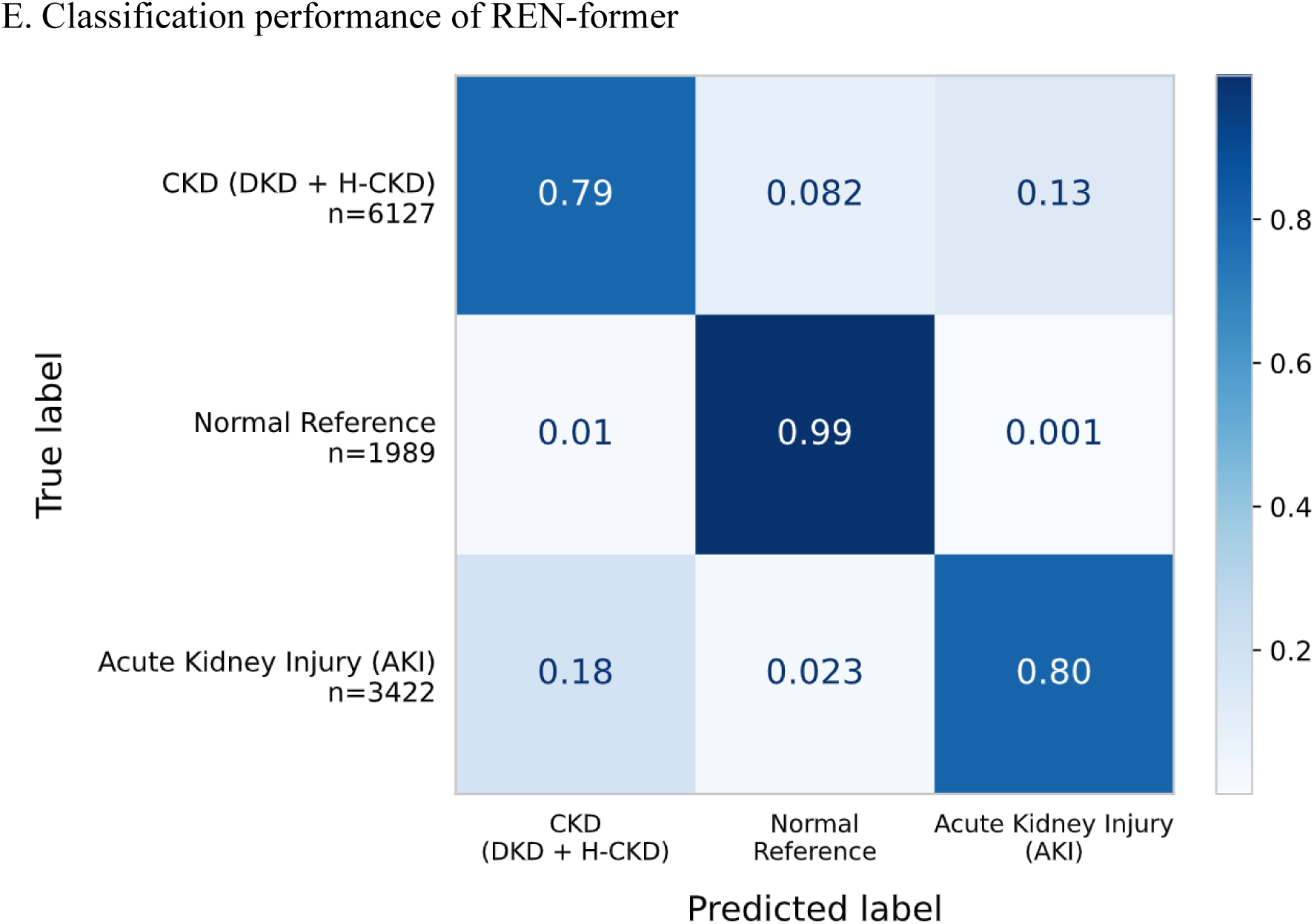
Construction and evaluation of REN-former (A) Overview of the REN-former analytical framework. Publicly available human kidney single-cell transcriptomes from Normal Reference, AKI, and CKD samples were used to fine-tune Geneformer. The resulting REN-former model was applied to in silico gene perturbation analysis, and selected candidate genes were further evaluated using summary-data-based Mendelian randomization and colocalization analyses. (B) Reference UMAP visualization of kidney cells colored by disease state. Diabetic kidney disease and hypertensive CKD samples were combined into the CKD category. (C) Reference UMAP visualization of kidney cells colored by the original kidney cell or structure annotation. (D) Heatmap of REN-former classification scores for cells in the test dataset. Rows represent individual cells and columns represent the three predicted disease states. Side and top annotations indicate the true and predicted class labels, respectively. (E) Row-normalized confusion matrix for cell-level classification in the held-out test set. Rows indicate true labels and columns indicate predicted labels. Values represent the proportion of cells within each true class assigned to each predicted class; therefore, diagonal values correspond to class-specific recall. Numbers in parentheses beside the row labels indicate the number of cells in each true class.

**Table 1.** Demographic and clinical characteristics of participants included in the GSE183276 dataset. Characteristics are presented according to the disease-state categories used for REN-former training and subsequent analyses. Values are shown as number (percentage) or mean, as indicated. Diabetic kidney disease and hypertensive CKD samples were combined into the CKD category. eGFR and albuminuria categories are presented for the CKD group. Albuminuria data were available for 10 of the 15 participants with CKD; percentages for albuminuria categories were calculated among participants with available data.

| Characteristic | Normal Reference<br>(n=18) | AKI (n=12) | CKD (n=15) |
| --- | --- | --- | --- |
| Male, n (%) | 7 (38.9) | 9 (75.0) | 5 (33.3) |
| Age, years, mean | 45.1 | 51.7 | 62.3 |
| White race, n (%) | 16 (88.9) | 6 (50.0) | 12 (80.0) |
| Diabetes, n (%) | 0 (0.0) | 5 (41.7) | 13 (86.7) |
| Diabetes duration, years, mean | — | — | 16 |
| Hypertension, n (%) | 0 (0.0) | 4 (33.3) | 14 (93.3) |
| Hypertension duration, years, mean | — | 2.5 | 14.8 |
| eGFR category, mL/min/1.73 m<sup>2</sup>, n (%) |  |  |  |
| > 60 | — | — | 4 (26.7) |
| 50–59 | — | — | 1 (6.7) |
| 40–49 | — | — | 5 (33.3) |
| 30–39 | — | — | 3 (20.0) |
| 20–29 | — | — | 2 (13.3) |
| Albuminuria category, mg, n (%) |  |  |  |
| < 30 | — | — | 1 (10.0) |
| 30–299 | — | — | 3 (30.0) |
| 300–499 | — | — | 0 (0.0) |
| 500–999 | — | — | 3 (30.0) |
| ≥1,000 | — | — | 3 (30.0) |
| Missing, n | — | — | 5 |

We next fine-tuned the pretrained Geneformer model to classify individual kidney-cell transcriptomes into Normal Reference, AKI, or CKD states and designated the resulting model REN-former. Samples were divided at the participant level into training, validation, and test sets containing 34, 5, and 6 participants, respectively. Cells from the same participants were not shared across data partitions. The predicted class-probability distributions showed that cells from each disease group generally received higher probabilities for their corresponding state than for the alternative states (Figure 1D). The row-normalized confusion matrix yielded class-specific recall values of 0.99, 0.80, and 0.79 for Normal Reference, AKI, and CKD, respectively (Figure 1E).

These findings indicate that fine-tuning Geneformer on human kidney single-cell transcriptomes generated a model capable of capturing disease-state-associated transcriptomic differences among Normal Reference, AKI, and CKD cells. This trained model was subsequently used for in silico perturbation analyses of directional transcriptomic state shifts.

### Directional deletion perturbation reveals broad regulatory remodeling during the AKI-to-CKD transition

To identify genes whose loss promoted directional transitions between kidney disease states, we performed single-gene deletion analysis in proximal-tubule cells and quantified the resulting shift toward each prespecified goal state (Figure 2A). Directional candidates were defined as genes whose deletion produced a positive shift toward the goal state with an FDR-adjusted *P* value below 0.05.

**Figure 2.**
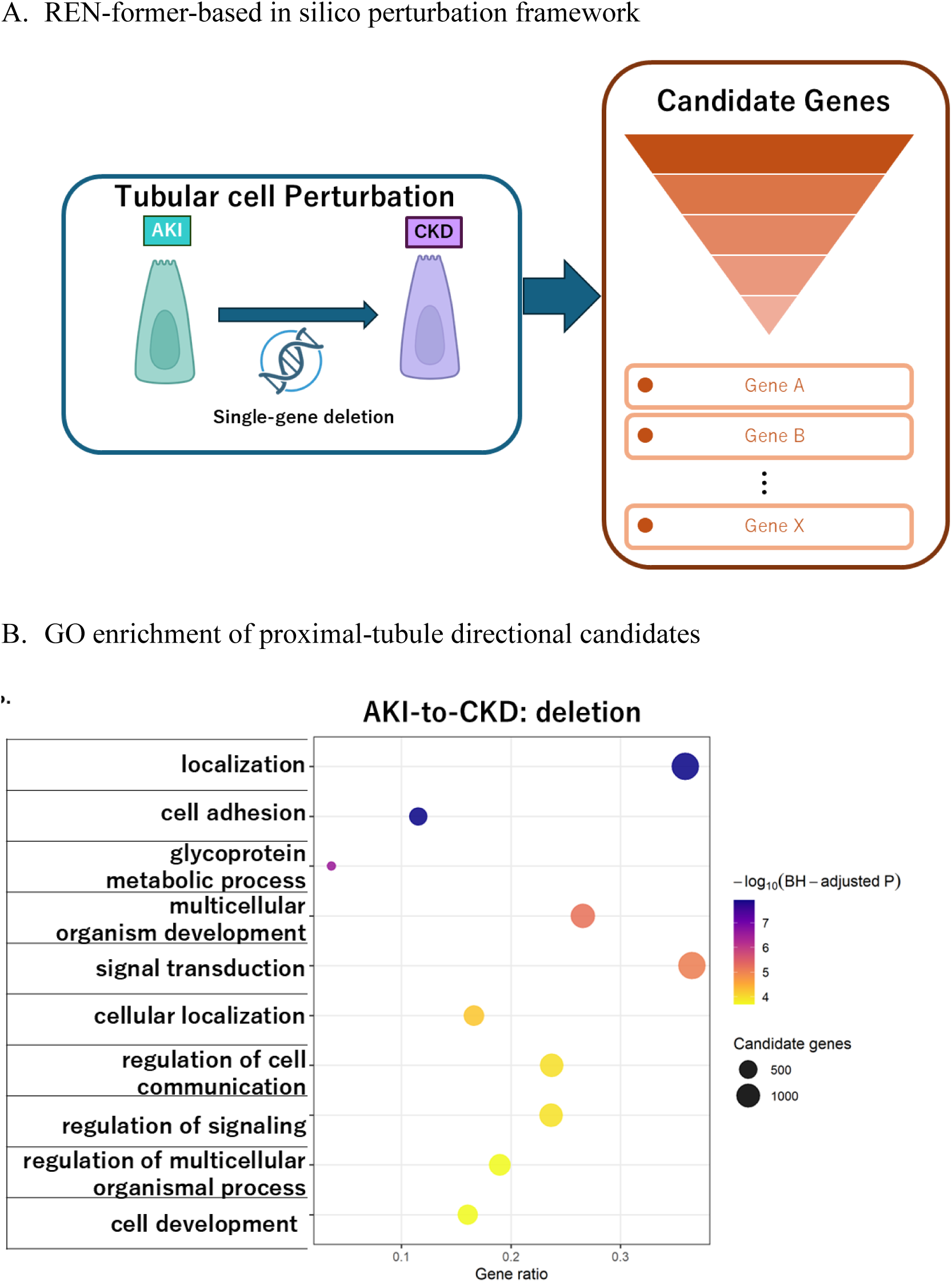
Directional deletion perturbation analysis identifies biological programs associated with the AKI-to-CKD transition. (A) Schematic overview of the in silico deletion analysis used to identify genes whose deletion shifted proximal-tubule transcriptomic states toward a specified goal state. Directional candidates were defined as genes producing a positive state shift with a false-discovery rate below 0.05. (B) Gene Ontology Biological Process enrichment of directional deletion candidates for the AKI-to-CKD transition in proximal-tubule cells. Enrichment analysis used all genes statistically evaluated in the corresponding perturbation analysis as the background. Redundant significant terms were reduced using Wang semantic similarity. The x-axis indicates the gene ratio, point size indicates the number of candidate genes contributing to each term, and point color indicates -log10 (Benjamini–Hochberg-adjusted *P* value). No GO Biological Process term was significantly enriched for the Normal Reference-to-AKI or Normal Reference-to-CKD deletion candidate sets after multiple-testing correction.

Application of this criterion to the proximal-tubule analysis identified 17 mapped candidates for the Normal Reference-to-AKI transition and 14 for the Normal Reference-to-CKD transition. Neither candidate set yielded a significantly enriched Gene Ontology Biological Process term after multiple-testing correction. In contrast, 4,456 mapped directional candidates were identified in proximal tubular cells for the AKI-to-CKD transition. Complete lists of deletion-derived directional candidate genes for all three transitions are provided in Supplementary Table 1. The corresponding all-cell analysis identified 1,420 candidates, of which 883 were shared with the proximal-tubule analysis (Supplementary Figure 1A). GO enrichment analysis of the 4,456 proximal-tubule candidates identified 63 significant nonredundant biological processes, indicating broad regulatory remodeling associated with a shift from the AKI-like to the CKD-like transcriptomic state (Figure 2B; Supplementary Table 2). The leading terms included localization, cell adhesion, glycoprotein metabolic process, multicellular organism development, signal transduction, cellular localization, and regulation of cell communication and signaling. Collectively, these results suggest that the AKI-to-CKD transition involves coordinated alterations in cellular organization, intercellular interactions, and developmental and signaling programs rather than a single dominant biological pathway.

### Directional overexpression analysis identifies mitochondrial, translational, and stress-response programs associated with kidney cell-state transitions

We next performed in silico single-gene overexpression analysis to identify genes whose increased activity was associated with directional shifts toward CKD-like transcriptomic states. Directional candidates were defined as genes whose overexpression produced a positive shift toward the prespecified goal state with an FDR-adjusted *P* value below 0.05. GO Biological Process enrichment analysis was performed separately for the Normal Reference-to-CKD and AKI-to-CKD transitions using the genes statistically evaluated in each perturbation analysis as the corresponding background.

For the Normal Reference-to-CKD transition, 330 mapped directional candidates were identified, yielding 139 significant nonredundant GO Biological Process terms after multiple-testing correction and semantic-similarity reduction (Figure 3A; Supplementary Table 3A and 4A). The leading enriched processes included cytoplasmic translation, oxidative phosphorylation, and processes related to cell fusion and differentiation. These findings indicate that increased activity of genes involved in protein synthesis, mitochondrial energy metabolism, and cellular differentiation was associated with a transcriptomic shift from the Normal Reference-like toward the CKD-like state.

**Figure 3.**
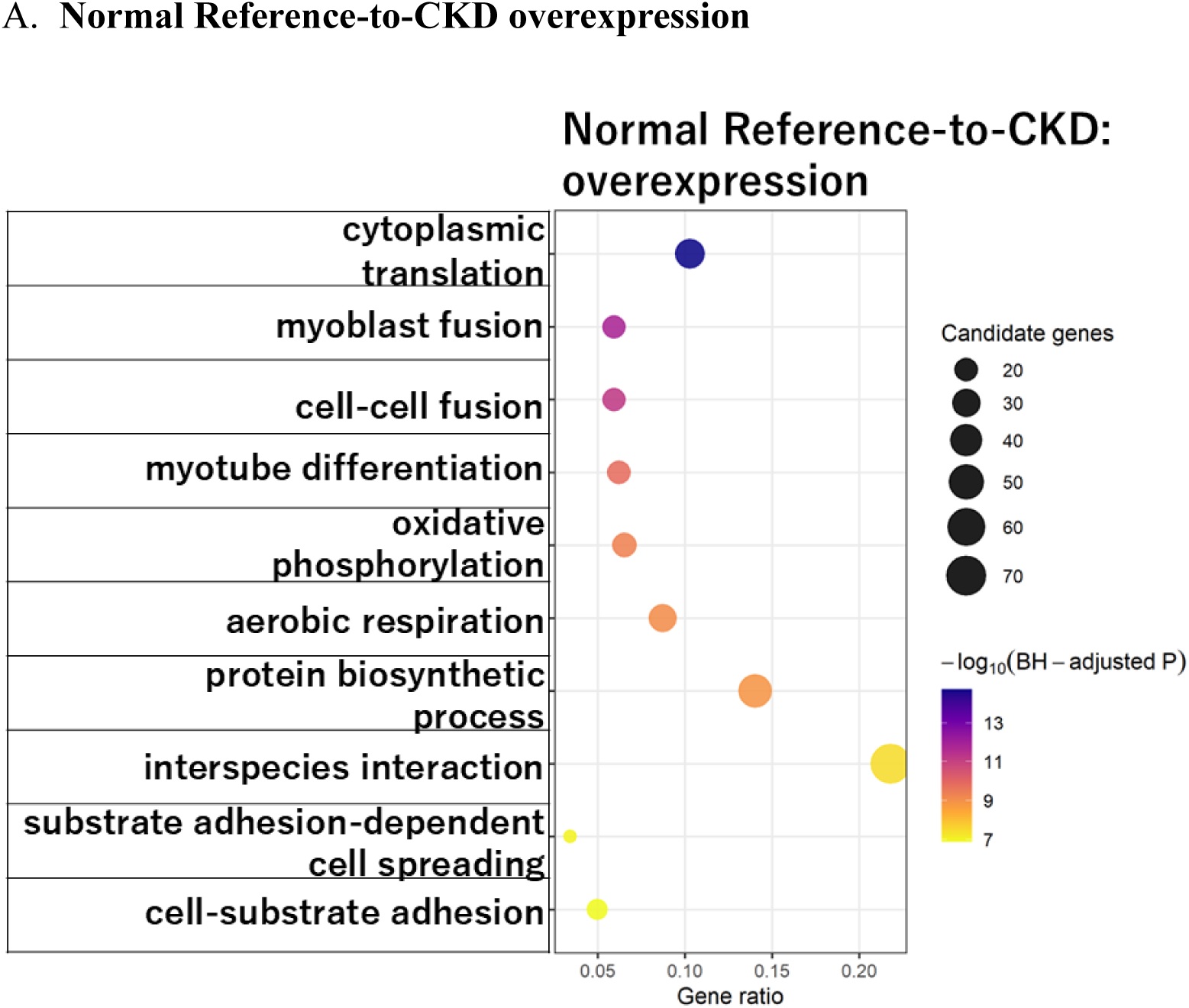
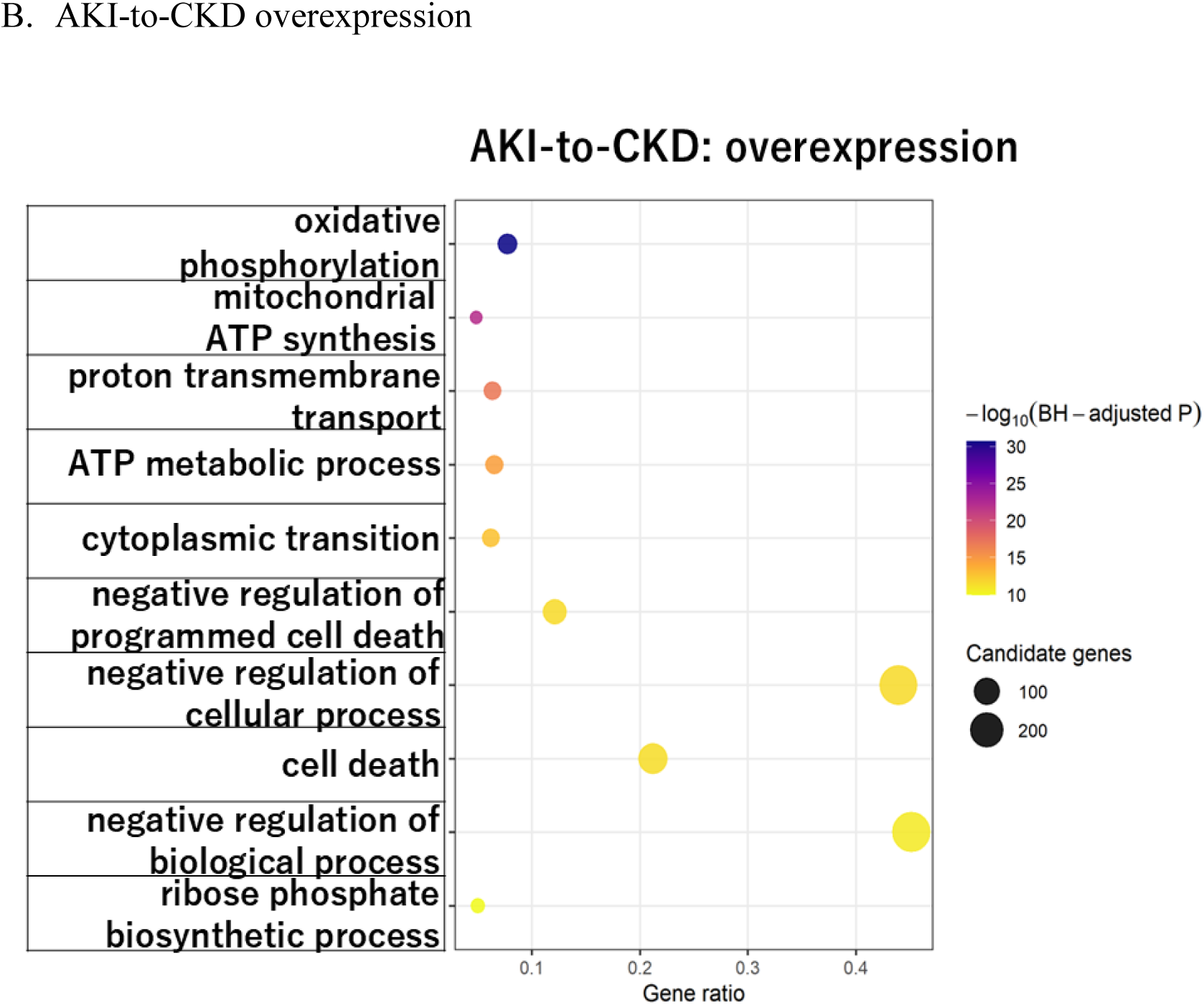
Gain-of-function perturbation identifies biological programs associated with transitions toward CKD-like states. (A) Gene Ontology (GO) Biological Process enrichment analysis of genes whose in silico overexpression shifted Normal Reference–like proximal tubular cells toward a CKD-like state. The representative nonredundant terms shown include cytoplasmic translation, cell– cell fusion and differentiation, oxidative phosphorylation, aerobic respiration, protein biosynthesis, and cell–substrate adhesion. (B) GO Biological Process enrichment analysis of genes whose in silico overexpression shifted AKI-like proximal tubular cells toward a CKD-like state. The representative nonredundant terms shown were dominated by mitochondrial energy metabolism and protein synthesis, including oxidative phosphorylation, proton motive force–driven mitochondrial ATP synthesis, proton transmembrane transport, ATP metabolic processes, and cytoplasmic translation, together with processes related to the regulation of cell death. Candidate genes were defined as genes whose overexpression induced a positive shift toward the specified goal state with a false discovery rate–adjusted P value <0.05. Enrichment analysis used all genes statistically evaluated in the corresponding perturbation analysis as the background. Redundant significant terms were reduced using Wang semantic similarity, and representative nonredundant terms are displayed. The x-axis indicates the gene ratio, point size indicates the number of candidate genes contributing to each term, and point color indicates -log10 of the Benjamini–Hochberg-adjusted P value. Complete lists of significant nonredundant GO Biological Process terms and their associated genes are provided in Supplementary Table 3. Abbreviations: AKI, acute kidney injury; CKD, chronic kidney disease; GO, Gene Ontology.

For the AKI-to-CKD transition, 685 mapped directional candidates were identified, yielding 136 significant nonredundant GO Biological Process terms (Figure 3B; Supplementary Table 3B and 4B). Prominent terms included oxidative phosphorylation, mitochondrial ATP synthesis coupled to electron transport, proton transmembrane transport, ATP metabolic processes, and cytoplasmic translation. Thus, the AKI-to-CKD shift induced by gene overexpression was strongly associated with coordinated changes in mitochondrial bioenergetics and protein synthesis.

Collectively, directional overexpression analysis identified mitochondrial and translational programs shared by transitions toward CKD-like states. These results suggest that the consequences of increasing individual gene activity depend on the broader transcriptomic context and that metabolic and biosynthetic processes may participate in remodeling of proximal tubular-cell states during CKD development.

### Reverse perturbation analysis identifies chemical-homeostasis and stress-response programs associated with shifts from CKD-like toward Normal Reference-like states

We next performed reverse perturbation analyses to identify genes whose deletion or overexpression shifted CKD-like proximal tubular cells toward the Normal Reference-like transcriptomic state. The deletion and overexpression analyses were evaluated separately using the same directional candidate criterion.

Deletion analysis identified 34 mapped directional candidates whose loss shifted CKD-like cells toward the Normal Reference-like state. GO enrichment analysis identified 38 significant nonredundant Biological Process terms (Figure 4A; Supplementary Table 5A and 6A). The leading terms included chemical homeostasis, cellular responses to chemical stimuli, and detoxification of copper, cadmium, and other inorganic compounds. These findings suggest that reduced activity of genes involved in metal handling and chemical-stress responses was associated with a shift away from the CKD-like transcriptomic state.

**Figure 4.**
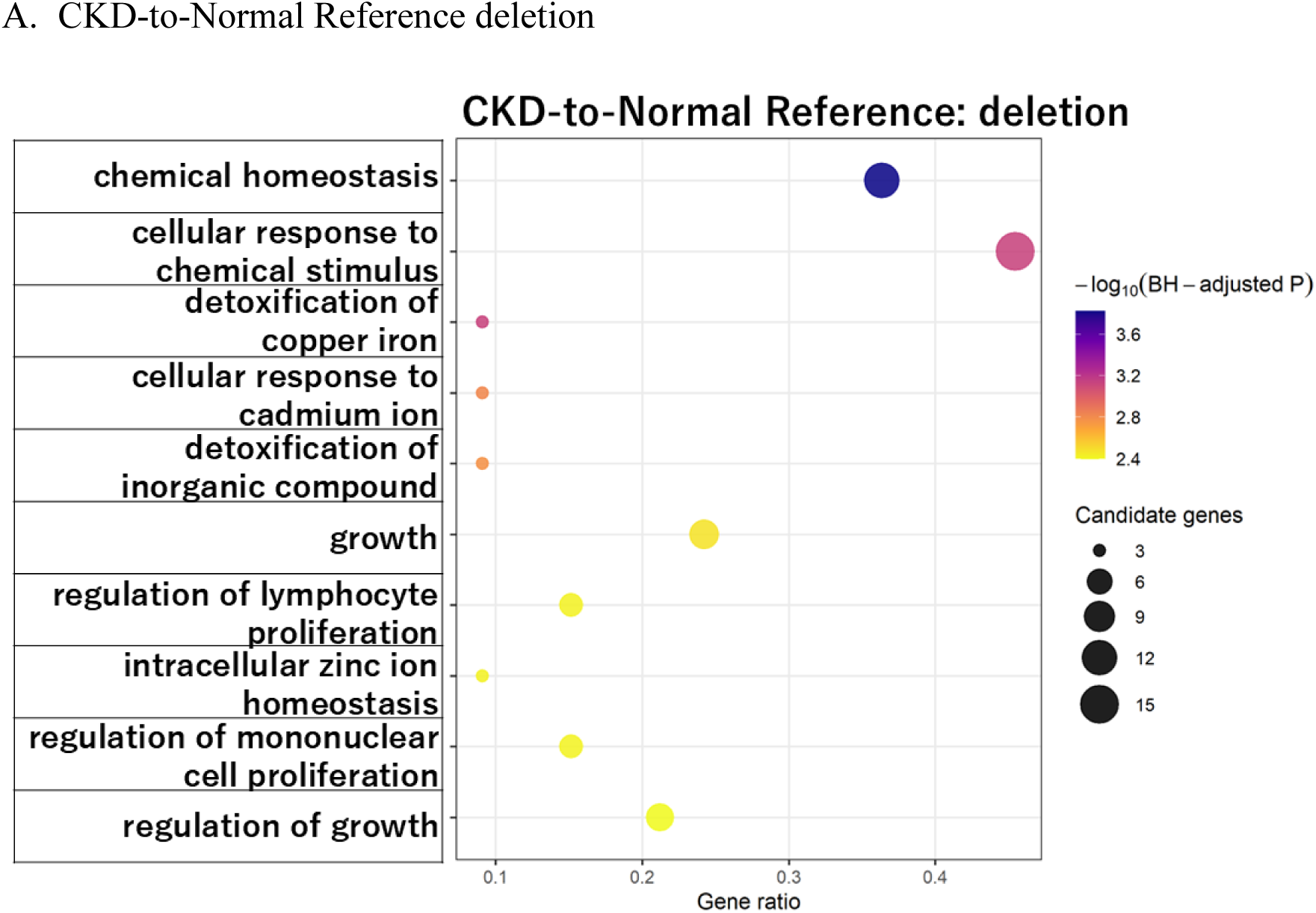
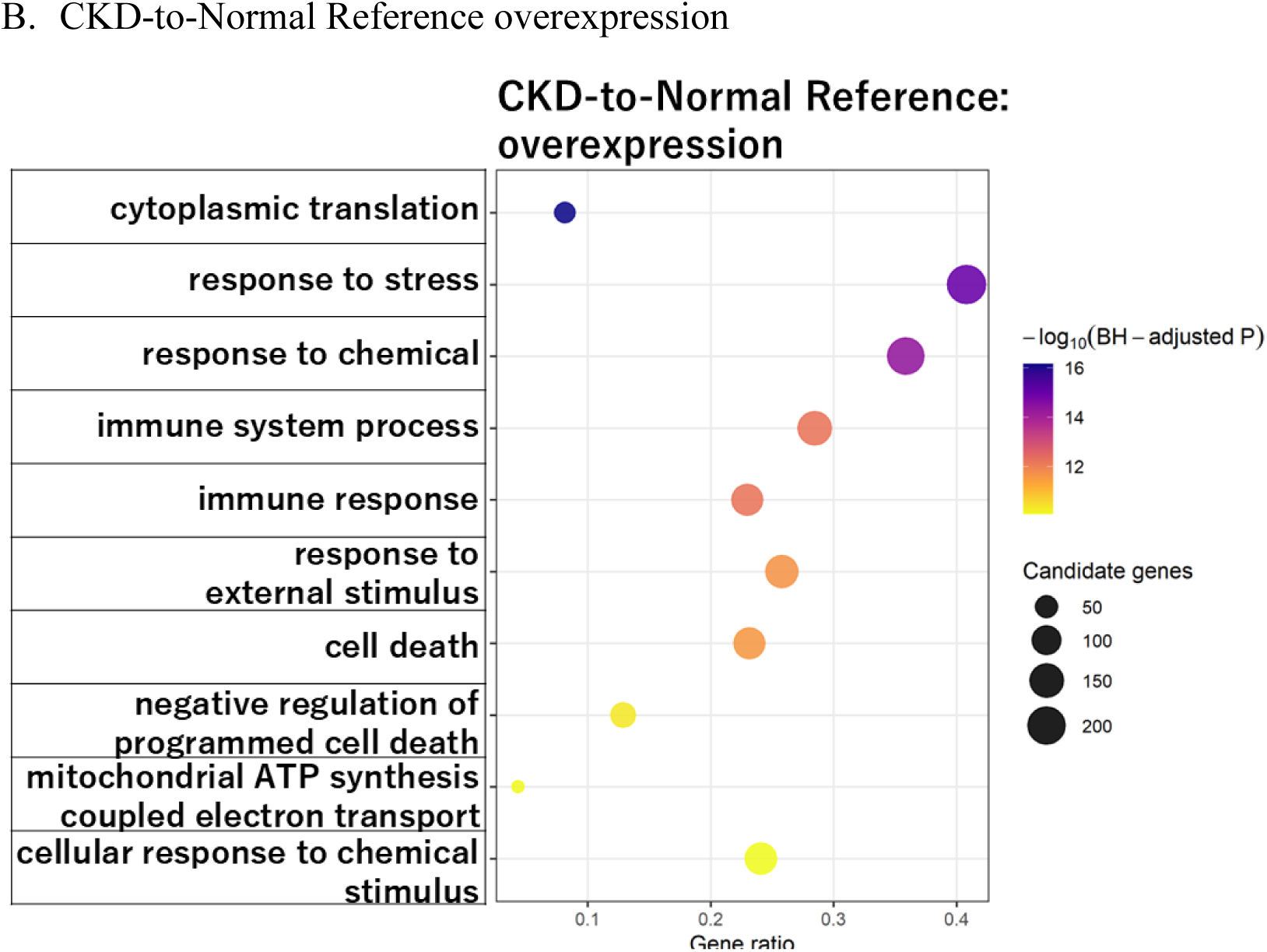
Reverse perturbation identifies biological programs associated with shifts from a CKD-like toward a Normal Reference–like state. (A) Gene Ontology (GO) Biological Process enrichment analysis of genes whose in silico deletion shifted CKD-like proximal tubular cells toward a Normal Reference–like state. The representative nonredundant terms shown include chemical homeostasis, cellular responses to chemical stimuli, detoxification of copper ions and inorganic compounds, cellular responses to cadmium ions, intracellular zinc-ion homeostasis, regulation of growth, and regulation of lymphocyte and mononuclear-cell proliferation. (B) GO Biological Process enrichment analysis of genes whose in silico overexpression shifted CKD-like proximal tubular cells toward a Normal Reference–like state. The representative nonredundant terms shown include cytoplasmic translation, responses to stress and chemical or external stimuli, immune-system processes and immune responses, regulation of cell death, mitochondrial ATP synthesis–coupled electron transport, and cellular responses to chemical stimuli. Candidate genes were defined as genes whose perturbation induced a positive shift toward the Normal Reference goal state with a false discovery rate–adjusted P value <0.05. Enrichment analysis used all genes statistically evaluated in the corresponding perturbation analysis as the background. Redundant significant terms were reduced using Wang semantic similarity, and representative nonredundant terms are displayed. The x-axis indicates the gene ratio, point size indicates the number of candidate genes contributing to each term, and point color indicates -log10 of the Benjamini–Hochberg-adjusted P value. Complete lists of significant nonredundant GO Biological Process terms and their associated genes are provided in Supplementary Table 5.

In contrast, overexpression analysis identified 558 mapped directional candidates associated with a shift from the CKD-like toward the Normal Reference-like state and yielded 143 significant nonredundant GO Biological Process terms (Figure 4B; Supplementary Table 5B and 6B). Prominent processes included cytoplasmic translation, responses to stress and chemical stimuli, immune-system processes, immune responses, and regulation of cell death. Thus, increased activity of genes involved in protein synthesis, stress adaptation, immune regulation, and cell-survival control was associated with movement toward the Normal Reference-like transcriptomic state.

Together, the reverse perturbation analyses identified distinct programs associated with shifts away from the CKD-like state. Gene deletion predominantly implicated metal and chemical homeostasis, whereas gene overexpression highlighted translational, stress-response, immune, and cell-death regulatory processes. These findings underscore that transcriptomic recovery from a CKD-like state may involve coordinated modulation of multiple cellular functions rather than reversal of a single disease-associated pathway.

### Conventional transcriptomic analysis identifies metabolic suppression and inflammatory activation in diseased proximal tubules

To compare REN-former with a conventional approach, we separately performed participant-level pseudobulk differential expression analysis and Hallmark gene set enrichment analysis (GSEA) using proximal tubular cells from the same GSE183276 dataset (Figure 5A). REN-former estimated how changing one gene shifted cells between states. In contrast, conventional analysis described the gene-expression differences that were already present between disease groups. The two approaches therefore provided different but complementary information about AKI and CKD.

**Figure 5.**
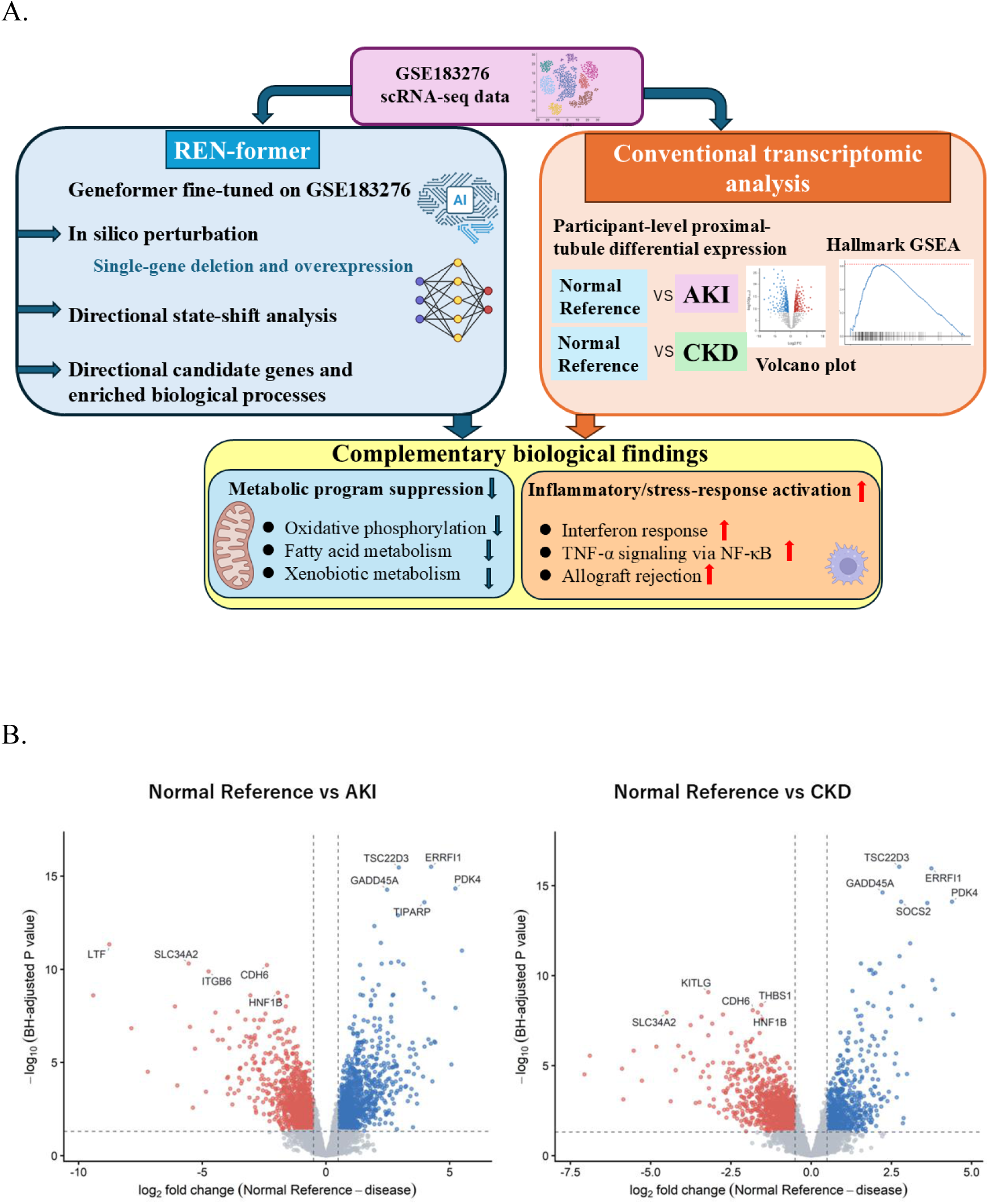
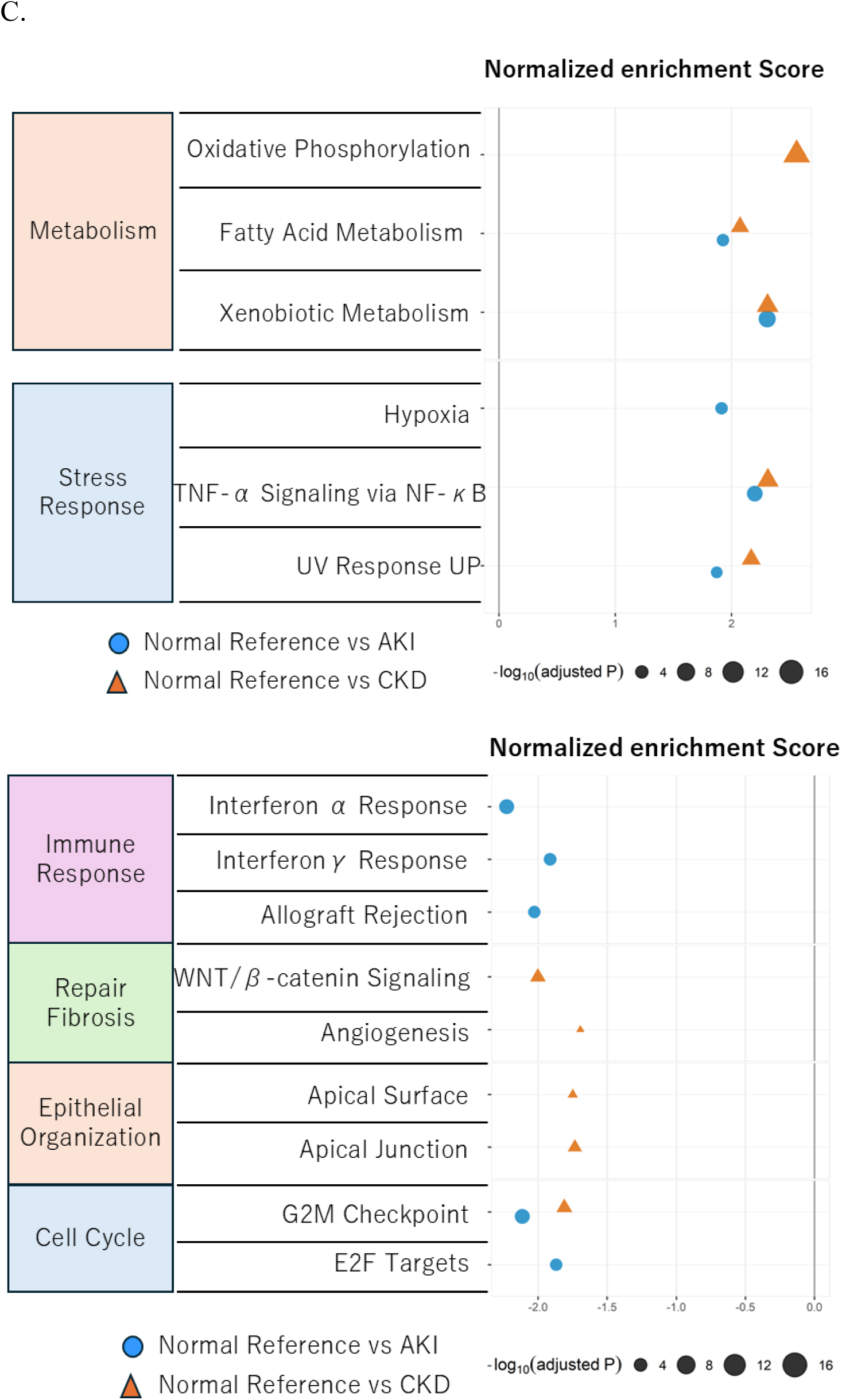
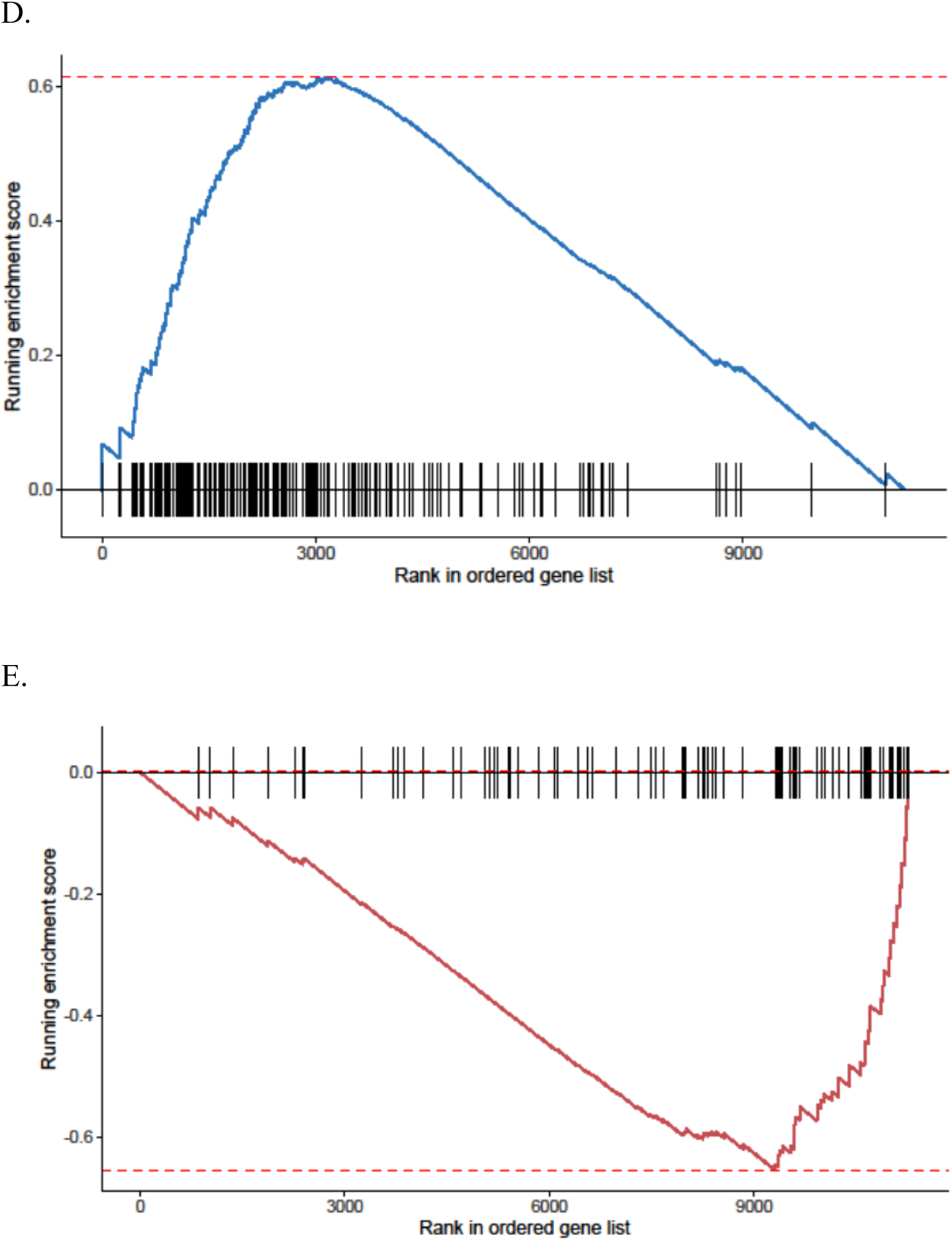
Integration of REN-former perturbation analysis with conventional transcriptomic analysis reveals metabolic suppression and inflammatory activation in diseased proximal tubules. (A) Schematic comparison of REN-former and conventional transcriptomic analyses using the GSE183276 scRNA-seq dataset. REN-former used modeled single-gene deletion and overexpression to prioritize genes and biological processes associated with directional transcriptomic state shifts. Conventional analysis used participant-level proximal-tubule pseudobulk profiles to compare Normal Reference with AKI and CKD, followed by differential expression analysis and Hallmark gene set enrichment analysis. The metabolic, inflammatory, and stress-response programs observed in the conventional analysis provided biological context for REN-former-prioritized processes but were not interpreted as direct validation of the predicted perturbation effects. (B) Volcano plots showing differential gene expression in proximal tubular cells for Normal Reference versus AKI and Normal Reference versus CKD comparisons. The x-axis represents the log2 fold change, and the y-axis represents -log10 of the false discovery rate (FDR). Blue points indicate genes expressed at significantly higher levels in Normal Reference proximal tubules, red points indicate genes expressed at significantly higher levels in AKI or CKD proximal tubules, and gray points indicate genes that did not meet the prespecified significance criteria. (C) Hallmark pathway enrichment in the Normal Reference-versus-AKI and Normal Reference-versus-CKD comparisons. Circles represent the Normal Reference-versus-AKI comparison, and triangles represent the Normal Reference-versus-CKD comparison. The x-axis shows the normalized enrichment score (NES); positive values indicate enrichment in Normal Reference proximal tubules, whereas negative values indicate enrichment in diseased proximal tubules. Symbol size represents -log10 of the Benjamini–Hochberg-adjusted *P* value. Among pathways with a Benjamini–Hochberg-adjusted P value <0.05, up to five pathways with the largest absolute NES were displayed separately for each contrast and enrichment direction. Metabolic pathways, including oxidative phosphorylation, fatty acid metabolism, and xenobiotic metabolism, were enriched toward Normal Reference. Interferon-response, allograft-rejection, WNT/β-catenin-signaling, epithelial-organization, and cell-cycle pathways showed enrichment toward diseased proximal tubules, whereas other stress-response pathways showed contrast- and pathway-dependent directions. (D) GSEA enrichment plot for the Hallmark oxidative phosphorylation pathway in the Normal Reference-versus-CKD comparison. Oxidative phosphorylation genes were enriched in Normal Reference proximal tubules (NES = 2.56; nominal *P* = 1.97 × 10⁻¹⁸; Benjamini– Hochberg-adjusted *P* = 9.84 × 10⁻¹⁷), consistent with suppression of mitochondrial energy metabolism in CKD. (E) GSEA enrichment plot for the Hallmark interferon-α response pathway in the Normal Reference-versus-AKI comparison. Interferon-α response genes were enriched in AKI proximal tubules (NES = -2.23; nominal *P* = 3.08 × 10⁻⁹; Benjamini–Hochberg-adjusted *P* = 3.85 × 10⁻⁸), indicating activation of inflammatory signaling during acute kidney injury.

Participant-level pseudobulk analysis revealed widespread transcriptional differences between Normal Reference and diseased proximal tubules (Figure 5B). Among 11,273 genes tested, 2,435 met the prespecified differential-expression criteria of a Benjamini–Hochberg false discovery rate (FDR) < 0.05 and an absolute log2 fold change ≥ 0.5 in the Normal Reference-versus-AKI comparison. Of these, 1,193 genes were expressed at higher levels in Normal Reference proximal tubules and 1,242 were expressed at higher levels in AKI proximal tubules. In the Normal Reference-versus-CKD comparison, 2,106 genes met the same criteria, including 790 genes expressed at higher levels in Normal Reference proximal tubules and 1,316 expressed at higher levels in CKD proximal tubules.

Pre-ranked Hallmark gene set enrichment analysis further identified coordinated pathway-level differences between Normal Reference and diseased proximal tubules (Figure 5C). Of the 50 Hallmark pathways tested, 27 were significantly enriched in the Normal Reference-versus-AKI comparison and 34 in the Normal Reference-versus-CKD comparison at BH-adjusted P < 0.05. In the Normal Reference-versus-AKI comparison, 17 pathways had positive normalized enrichment scores (NES), indicating enrichment toward Normal Reference, whereas 10 had negative NES, indicating enrichment toward AKI. In the Normal Reference-versus-CKD comparison, 20 pathways were enriched toward Normal Reference and 14 toward CKD. For visualization, Figure 5C displays up to the five pathways with the strongest absolute NES in each direction and comparison among pathways meeting BH-adjusted P < 0.05. The enriched pathways encompassed metabolic, stress-response, immune, repair/fibrosis, epithelial-organization, and cell-cycle programs.

Oxidative phosphorylation showed particularly strong enrichment in Normal Reference proximal tubules relative to CKD proximal tubules (NES = 2.56; nominal P = 1.97 × 10⁻¹⁸; BH-adjusted P = 9.84 × 10⁻¹⁷) (Figure 5D). The concentration of oxidative-phosphorylation genes toward the Normal Reference end of the ranked gene list was consistent with reduced mitochondrial energy-metabolism programs in CKD proximal tubules.

Conversely, the interferon-α response pathway was enriched toward AKI proximal tubules in the Normal Reference-versus-AKI comparison (NES = -2.23; nominal P = 3.08 × 10⁻⁹; BH-adjusted P = 3.85 × 10⁻⁸) (Figure 5E). The negative enrichment score and concentration of interferon-response genes toward the AKI end of the ranked list were consistent with increased interferon-associated inflammatory signaling during AKI.

Conventional transcriptomic analyses provided biological context for the processes prioritized by REN-former. Diseased proximal tubules showed suppression of metabolic programs, including oxidative phosphorylation, fatty acid metabolism, and xenobiotic metabolism, together with activation of inflammatory and stress-response programs, including interferon responses and TNF-α signaling via NF-κB. These observed disease-associated programs were broadly related to processes highlighted by REN-former but did not constitute direct validation of the predicted perturbation effects. The two approaches addressed distinct but complementary questions: conventional analysis characterized expression differences already present between disease groups, whereas REN-former estimated directional transcriptomic shifts following modeled single-gene perturbation.

### Human genetic analyses support kidney function associations of REN-former-prioritized genes

To investigate whether genes prioritized by REN-former were supported by human kidney genetic data, we integrated the perturbation-derived candidates with tubular expression quantitative trait locus (eQTL) and estimated glomerular filtration rate (eGFR) genome-wide association study (GWAS) summary statistics (Figure 6A). Genes identified in four perturbation settings—Normal Reference-to-CKD deletion, Normal Reference-to-CKD overexpression, CKD-to-Normal Reference deletion, and CKD-to-Normal Reference overexpression—were included. After removing duplicated genes and genes located within the major histocompatibility complex region, 199 unique REN-former-prioritized genes were evaluated using summary-data-based Mendelian randomization (SMR).

**Figure 6.**
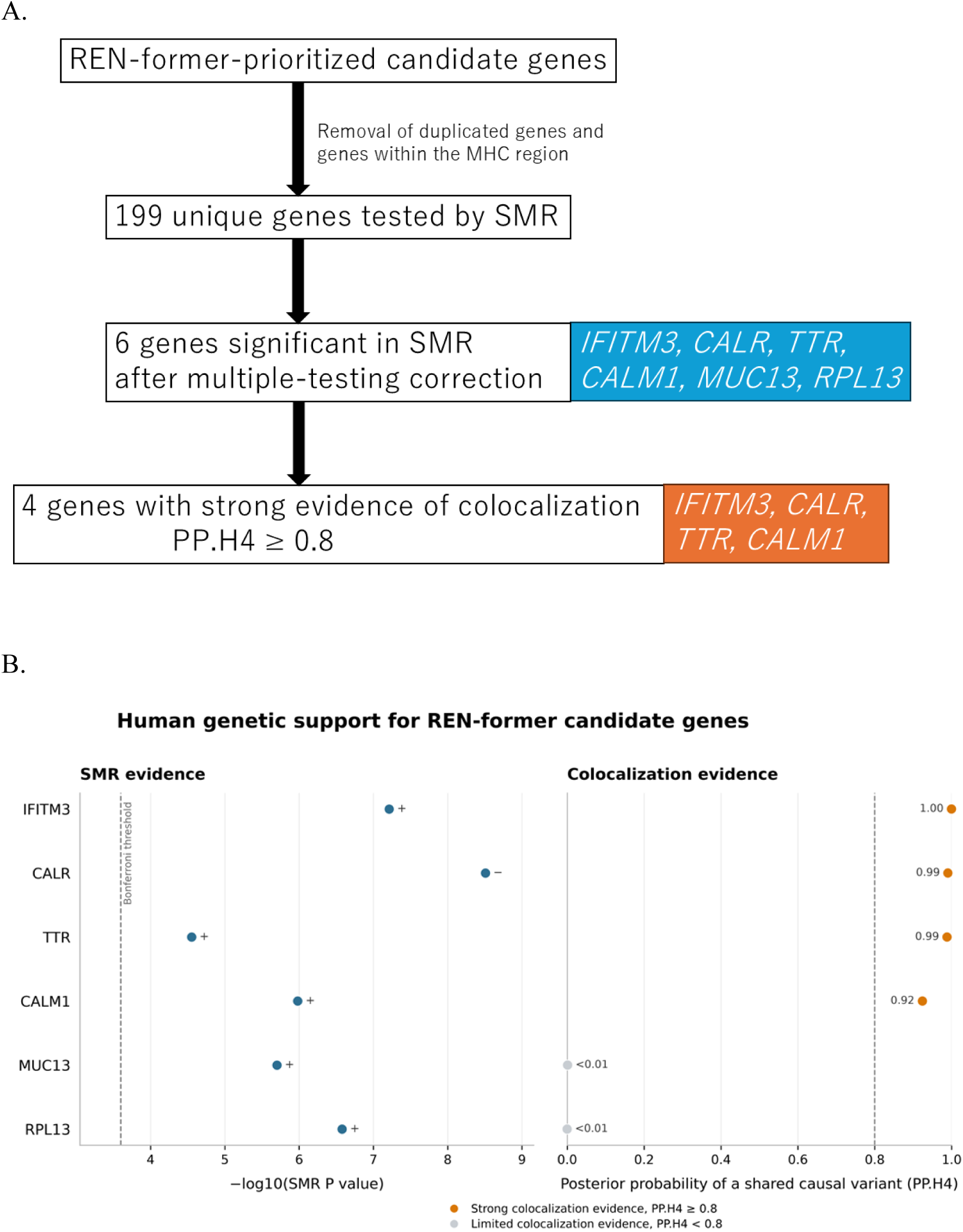
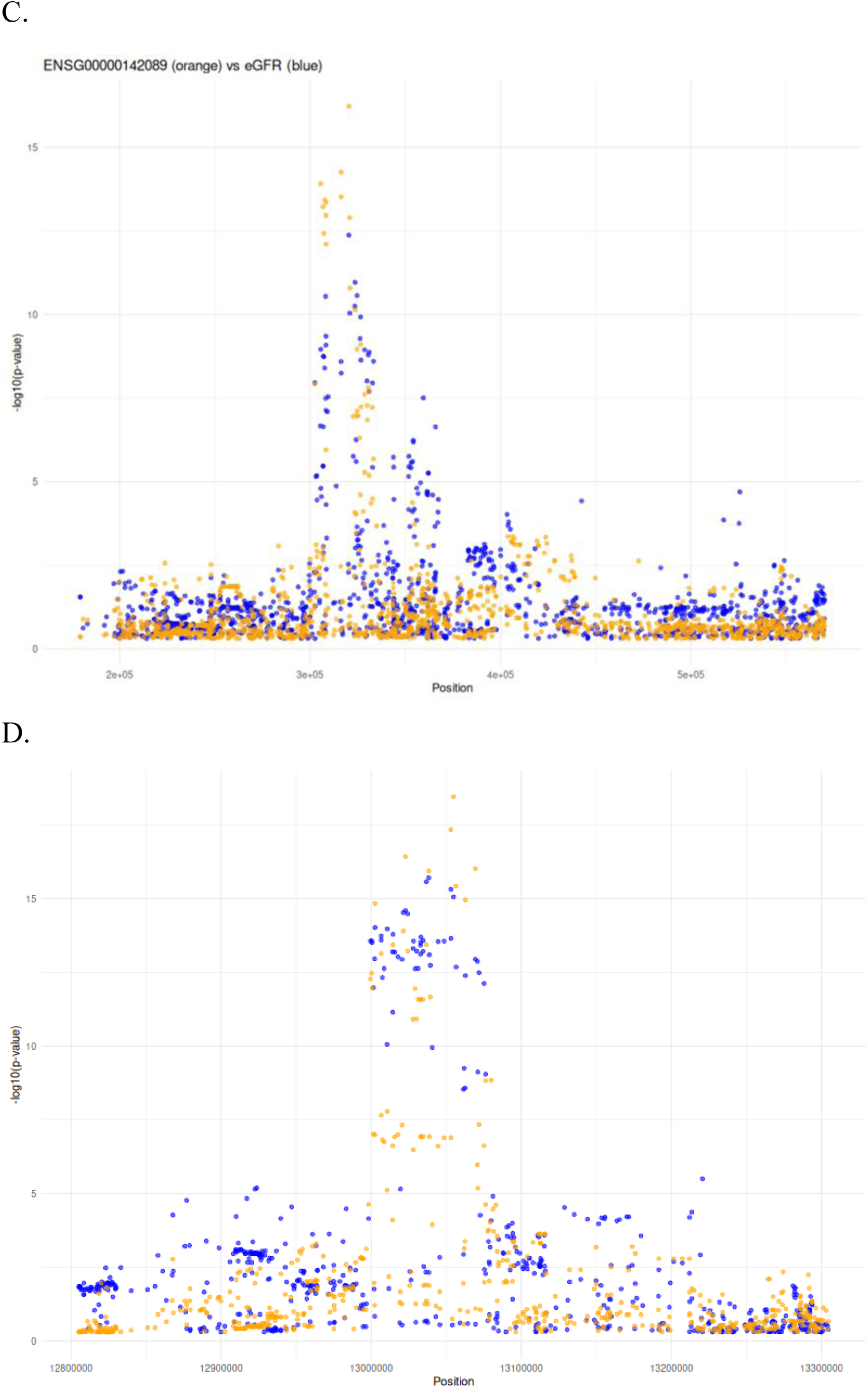
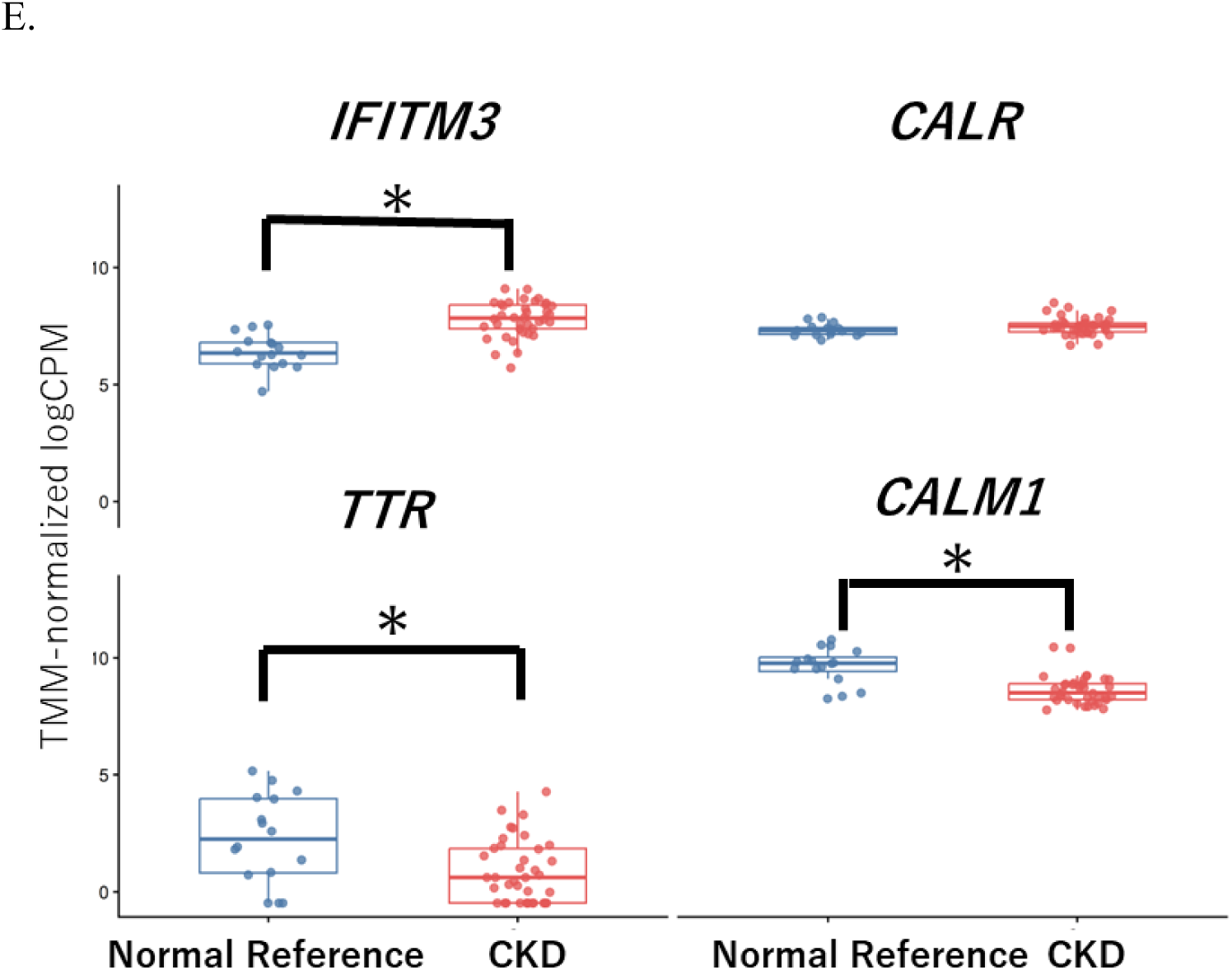
Human genetic analyses support kidney function associations of REN-former–prioritized genes. (A) Workflow for the integration of REN-former–prioritized candidate genes with human kidney genetic data. After removal of duplicated genes and genes located within the major histocompatibility complex region, 199 unique candidate genes were evaluated by summary-data-based Mendelian randomization (SMR) analysis using tubular expression quantitative trait locus (eQTL) and estimated glomerular filtration rate (eGFR) genome-wide association study (GWAS) summary statistics. Six genes—*IFITM3, CALR, TTR, CALM1, MUC13*, and *RPL13*— met the Bonferroni-corrected SMR threshold, passed the HEIDI test, and showed SMR effect directions concordant with the corresponding REN-former predictions. Subsequent colocalization analysis provided strong evidence consistent with a shared causal variant for *IFITM3, CALR, TTR*, and *CALM1*, with posterior probabilities for hypothesis 4 (PP.H4) ≥0.8. (B) Summary of the SMR and colocalization results for the six genes selected for colocalization analysis based on the prespecified SMR, HEIDI, and directional-concordance criteria. The left panel shows -log10-transformed SMR *P* values. The vertical dashed line indicates the Bonferroni-corrected significance threshold, and the plus and minus signs indicate the direction of the SMR effect. No evidence of heterogeneity was detected by the heterogeneity in dependent instruments (HEIDI) test for any of the displayed genes (HEIDI *P* >0.05). The right panel shows the posterior probability that the tubulointerstitial eQTL and eGFR GWAS associations share a causal variant (PP.H4). Orange points indicate strong colocalization evidence (PP.H4 ≥0.8), whereas gray points indicate limited evidence of colocalization (PP.H4 <0.8). (C) Regional association plot for *IFITM3* showing the tubulointerstitial eQTL association signal for *IFITM3* expression and the eGFR GWAS association signal across the genomic region. Orange points represent the tubulointerstitial eQTL associations, and blue points represent the eGFR GWAS associations. The overlapping association patterns were consistent with a shared underlying genetic signal influencing both *IFITM3* expression and kidney function. (D) Regional association plot for *CALR* showing the tubular eQTL association signal for *CALR* expression and the eGFR GWAS association signal across the genomic region. Orange points represent the tubular eQTL associations, and blue points represent the eGFR GWAS associations. The overlapping association patterns support the presence of a shared causal variant influencing both *CALR* expression and kidney function. (E) Participant-level expression of the four colocalized genes in proximal tubular cells from expanded-KPMP participants not overlapping GSE183276. The primary scRNA-seq analysis included 16 Normal Reference and 38 CKD participants. *IFITM3* was higher in CKD, whereas *TTR* and *CALM1* were lower in CKD. *CALR* did not differ significantly between the groups. Each point represents one participant, and boxes indicate the median and interquartile range. The observed expression directions were concordant with REN-former predictions for *CALR, TTR*, and *CALM1*, although *CALR* was not significant, whereas *IFITM3* showed the opposite direction.

Six genes—*IFITM3, CALR, TTR, CALM1, MUC13*, and *RPL13*— met the prespecified SMR and HEIDI criteria and showed SMR effect directions concordant with the corresponding REN-former predictions. These genes were taken forward for colocalization analysis (Figure 6B; Table 2). No significant heterogeneity was detected by the heterogeneity in dependent instruments test for these genes (HEIDI P > 0.05). Five of the six genes— *IFITM3, TTR, CALM1, MUC13,* and *RPL13*—were derived from the CKD-to-Normal Reference overexpression analysis, in which increased gene activity was associated with a transcriptomic shift toward a Normal Reference-like state. In contrast, *CALR* was identified in the Normal Reference-to-CKD overexpression analysis, in which increased expression was associated with a shift toward a CKD-like state. Thus, the human genetic analysis provided support for both recovery-associated and disease-promoting regulators identified by REN-former.

**Table 2.**
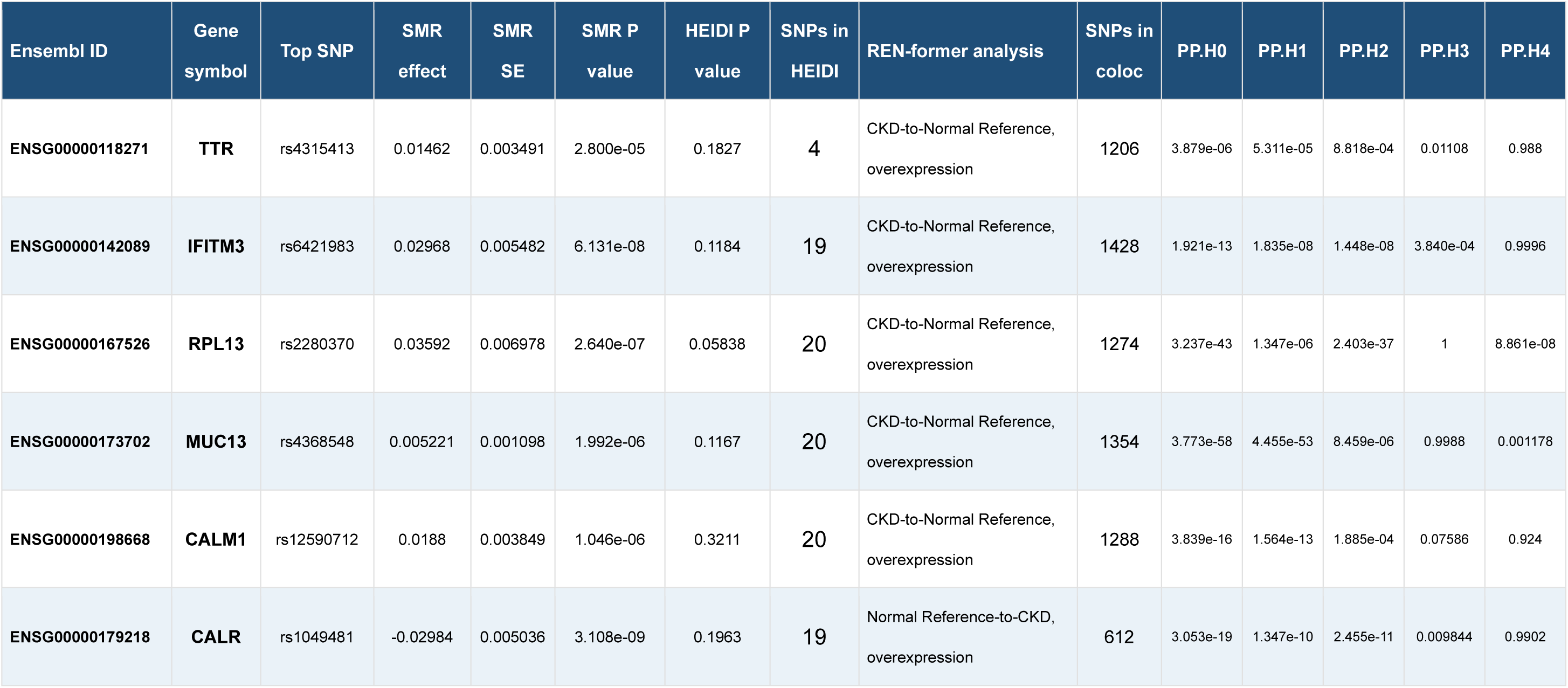
Summary-data-based Mendelian randomization and colocalization results for REN-former-prioritized genes. SMR and colocalization results for the six REN-former-prioritized genes that met the Bonferroni-corrected SMR threshold, showed no significant heterogeneity by the HEIDI test, and had SMR effect directions concordant with the corresponding REN-former-predicted perturbation directions. The heterogeneity in dependent instruments test was used to assess heterogeneity of the SMR associations. Colocalization results are presented as the posterior probability for hypothesis 4 (PP.H4), representing the probability that the tubular expression quantitative trait locus and estimated glomerular filtration rate genome-wide association study signals share an underlying causal variant. Strong evidence of colocalization was defined as PP.H4 ≥0.8.

We next performed colocalization analysis to determine whether the tubulointerstitial eQTL and eGFR GWAS associations were consistent with a shared underlying genetic signal. Four genes—*IFITM3, CALR, TTR*, and *CALM1*—showed strong evidence of colocalization, defined as a posterior probability for hypothesis 4 (PP.H4) ≥ 0.8 (Figure 6B; Table 2). Regional association plots for *IFITM3* and *CALR* showed overlapping tubulointerstitial eQTL and eGFR GWAS association patterns across their respective genomic regions (Figure 6C, D). Similar regional association patterns were observed for *TTR* and *CALM1,* whereas *MUC13* and *RPL13* showed more limited evidence of colocalization (Supplementary Figure 2A–D).

We also examined the expression of the four colocalized genes using the participant-level proximal-tubule analysis. *CALM1* expression was significantly lower in AKI and CKD than in Normal Reference samples (Normal Reference versus AKI: log2 fold change = 0.78, FDR = 0.00061; Normal Reference versus CKD: log2 fold change = 0.60, FDR = 0.0034) (Supplementary Figure 3). *IFITM3, CALR,* and *TTR* did not meet the prespecified differential-expression criteria. Thus, three of the four genetically supported candidates were not identified as conventional differentially expressed genes.

We next evaluated the four colocalized genes in expanded KPMP data after excluding participants included or potentially included in GSE183276. The primary scRNA-seq analysis included 16 Normal Reference and 38 CKD participants. *IFITM3* was higher in CKD (log2 fold change = 1.39, four-gene FDR = 9.67 × 10⁻⁷), whereas *TTR* (log2 fold change = -1.90, four-gene FDR = 1.17 × 10⁻⁴) and *CALM1* (log2 fold change = -1.08, four-gene FDR = 6.64 × 10⁻⁶) were lower in CKD. *CALR* was not significantly different (Figure 6E). The expression changes of *TTR* and *CALM1* were concordant with the REN-former-predicted direction. *CALR* showed a concordant but nonsignificant trend, whereas *IFITM3* was significantly increased in the opposite direction. In the snRNA-seq replication analysis of 84 Normal Reference and 80 CKD participants, *CALM1* remained significantly lower in CKD (log2 fold change = -0.45, four-gene FDR = 1.68 × 10⁻⁵), whereas the other three genes did not meet the four-gene FDR threshold (Supplementary Figure 4).

Collectively, REN-former perturbation analysis, human genetic evidence, and expression analysis in non-overlapping KPMP participants prioritized *IFITM3, CALR, TTR,* and *CALM1* for further study. REN-former predictions and the gene expression changes observed in patients with CKD were concordant for some genes but not for others. In addition, the observed expression changes were not consistently reproduced in both the scRNA-seq and snRNA-seq analyses. These findings indicate that REN-former predictions, genetic analyses, and observed gene expression evaluate candidate genes from different perspectives and that integrating these complementary sources of evidence may help prioritize candidate genes.

## Discussion

In this study, we developed REN-former by fine-tuning Geneformer with human kidney single-cell transcriptomes from Normal Reference, AKI, and CKD samples. REN-former was used to predict how deletion or overexpression of individual genes might shift proximal tubular cells between these transcriptomic states. The analysis identified candidate genes and biological processes associated with kidney injury, progression toward CKD, and shifts from a CKD-like toward a Normal Reference-like state.

Geneformer was pretrained on a large collection of human single-cell transcriptomes and can be adapted to specific biological questions.^22^ Fine-tuning the model with kidney data allowed REN-former to learn features related to kidney cell types and disease states. This is important because the human kidney contains many cell types, and injured cells can show different transcriptional states within the same cell population.^17^ Candidate genes differed between the analyses of all kidney cells and proximal tubular cells, particularly for the AKI-to-CKD transition. This suggests that focusing on the relevant cell type may reveal cell-specific gene relationships. However, REN-former is a hypothesis-generating tool and does not replace experimental validation or necessarily outperform simpler methods.^35^

Deletion perturbation of AKI-like proximal tubular cells identified a much broader set of candidates associated with a shift toward a CKD-like state. These candidates were enriched in processes related to cellular localization, adhesion, intercellular communication, development, and signal transduction. These processes may contribute to changes in tubular organization and function during the transition toward a CKD-like state. Previous studies have demonstrated that mitochondrial dysfunction, defective fatty acid oxidation, oxidative stress, and ferroptotic stress contribute to proximal tubular injury and maladaptive kidney repair^13, 36^. Single-cell studies of AKI have also identified dynamic proximal tubular states characterized by metabolic suppression, oxidative stress, and activation of injury-response pathways^12, 15^. Recent evidence further indicates that sustained *VCAM1* expression in injured proximal tubular cells promotes immune-cell adhesion and tubular–immune crosstalk, potentially reinforcing failed repair during the AKI-to-CKD transition.^37^ Other experimental and single-cell multiomic studies have similarly shown that injury severity and persistent inflammatory programs influence whether tubular cells undergo successful or failed repair.^38, 39^

Overexpression perturbation of Normal Reference–like proximal tubular cells toward a CKD-like state was associated with cytoplasmic translation, oxidative phosphorylation, aerobic respiration, protein biosynthesis, cell–cell fusion or differentiation, and cell–substrate adhesion. More generally, these results suggest that altered coordination of protein synthesis, energy metabolism, cellular differentiation, and adhesion may accompany departure from a Normal Reference–like tubular state. Because the direction of an embedding shift does not specify whether a pathway is physiologically adaptive or harmful, individual candidates within these categories require functional evaluation before they can be considered disease-promoting regulators.

Reverse perturbation analysis identified distinct processes associated with shifts from a CKD-like toward a Normal Reference–like state. Deletion candidates were mainly related to chemical and metal-ion homeostasis, whereas overexpression candidates were related to stress responses, mitochondrial function, protein synthesis, immune regulation, and cell death. These findings suggest that departure from a CKD-like state may involve several cellular processes. However, the predicted effects may depend on the injury stage and cellular environment and should not be interpreted as direct therapeutic effects. Studies following kidney injury over time have also shown that recovery and CKD progression vary between kidney regions and over time.^40, 41^

The conventional gene-expression and Hallmark pathway analyses provided complementary information. Diseased proximal tubular cells showed reduced oxidative phosphorylation, fatty acid metabolism, and xenobiotic metabolism programs, together with increased interferon-associated inflammatory signaling and broader alterations in stress-response and tissue-remodeling programs. These findings are consistent with previous studies of impaired mitochondrial metabolism and persistent inflammation in injured proximal tubular cells. ^17, 36, 39^ Conventional analysis describes gene-expression differences already present between disease groups, whereas REN-former predicts how changing one gene may shift the transcriptomic state.

Integration with human kidney genetic data provided an additional layer of support for REN-former-prioritized candidates. Of the 199 REN-former-prioritized genes, six— *IFITM3, CALR, TTR, CALM1, MUC13,* and *RPL13*—met the prespecified SMR and HEIDI criteria and showed SMR effect directions concordant with the corresponding REN-former-predicted perturbation directions (Figure 6B; Table 2). Complete SMR results are provided in Supplementary Table 7. Colocalization analysis further identified strong evidence that the tubulointerstitial eQTL and eGFR GWAS signals shared an underlying genetic signal for *IFITM3, CALR, TTR*, and *CALM1*. Compartment-specific kidney eQTL analyses have demonstrated that integration of kidney gene-expression regulation with GWAS signals can improve the prioritization of genes and cell types underlying kidney function and disease.^42, 43^ *IFITM3* links the genetic results to the interferon-response program identified in diseased proximal tubules, whereas *CALR* provides a potential connection to endoplasmic reticulum proteostasis and calcium homeostasis. *CALM1* encodes calmodulin, a major calcium-binding protein that regulates calcium-dependent signaling. Increased urinary exosomal *CALM1* protein has previously been reported in diabetic kidney disease.^44^ In contrast, we observed lower intracellular *CALM1* mRNA expression in CKD proximal tubules. The role of *TTR* in acquired tubular injury and recovery is less well defined and may represent a relatively novel finding of the present analysis.

The higher *IFITM3* expression observed in CKD, despite the REN-former-predicted shift toward a Normal Reference-like state, may reflect a compensatory response or disease-associated interferon activation rather than a disease-promoting effect of *IFITM3* itself. This discordance also illustrates the distinction between cross-sectional disease-associated expression and the predicted effect of perturbing an individual gene.

Importantly, the primary aim of this study was to establish a framework for prioritizing candidate genes by integrating foundation-model predictions with human genetic and transcriptomic evidence, rather than to validate the biological function of a specific gene. The four colocalized genes showed different relationships between predicted perturbation effects and observed CKD-associated expression. *TTR* and *CALM1* showed significant expression changes concordant with the REN-former-predicted direction, *CALR* showed a concordant but nonsignificant trend, and *IFITM3* changed significantly in the opposite direction. These differences illustrate that REN-former predictions, genetic associations, and disease-associated expression provide distinct and complementary information. *CALM1* should therefore be considered one of several candidates that may be related to reduced eGFR and CKD-associated tubular states, requiring further experimental investigation.

## Limitations

This study has several limitations. REN-former was developed using one publicly available, cross-sectional dataset with a limited number of participants and heterogeneous AKI and CKD etiologies. The AKI and CKD samples were obtained from different participants, and longitudinal kidney outcomes after AKI were not available. Therefore, the predicted transitions represent shifts between transcriptomic states rather than changes observed over time in the same participants or cells. In silico deletion and overexpression modify model representations and cannot fully reproduce biological gene perturbation. Similarly, SMR and colocalization support gene prioritization but do not prove causality. The candidate genes should therefore be tested in experimental systems and independent human kidney datasets. The additional expression cohort was drawn from the same KPMP research consortium and therefore was not fully independent of the training dataset used to develop REN-former.

## Conclusion

REN-former provides a kidney-contextualized framework for generating hypotheses about genes and biological processes associated with directional tubular-cell state shifts. The results suggest that transitions among Normal Reference–like, AKI-like, and CKD-like states involve coordinated changes in epithelial organization, mitochondrial and translational activity, stress responses, immune signaling, and cell survival. Integration with human kidney genetic data further prioritized *IFITM3, CALR, TTR,* and *CALM1* as candidates for mechanistic investigation.

## Author contributions

IM conceived and designed the study. IM and SH performed data analysis, deep learning, and visualization. YH performed SMR and colocalization analyses. IM, SH, and YH wrote the manuscript. IM, SH, YH, TK, MI, SK, YK, and TT reviewed and edited the manuscript. NT and MN supervised the study.

## Declaration of generative AI and AI-assisted technologies in the writing process

During the preparation of this work, the authors used ChatGPT to assist with English language editing and organization of the manuscript text. The authors reviewed and edited the content and take full responsibility for the final version of the manuscript.

---

**Supplementary Figure 1.**
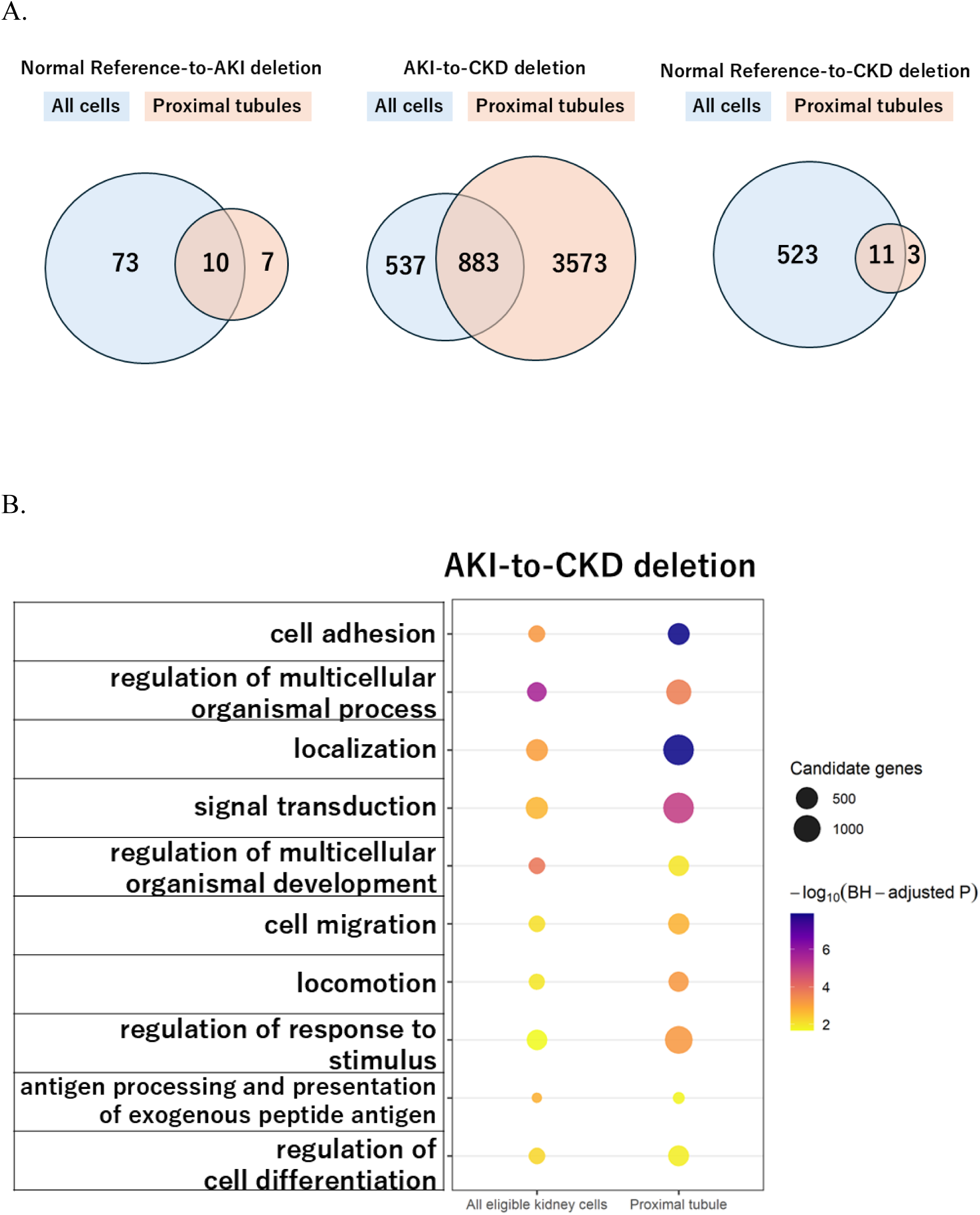
Comparison of deletion perturbation results obtained using all kidney cells and proximal tubule–restricted cells. (A) Venn diagrams show the overlap between significant candidate genes identified by deletion perturbation analysis using all eligible kidney cells and analysis restricted to proximal tubular cells. Results are presented for the Normal Reference-to-AKI, AKI-to-CKD, and Normal Reference-to-CKD state transitions. Candidate genes were defined as genes whose deletion induced a positive shift toward the predefined goal state, with a false discovery rate (FDR)-adjusted *P* value <0.05, and that were successfully mapped to Entrez Gene identifiers for downstream Gene Ontology enrichment analysis. Blue circles represent candidates identified using all eligible kidney cells, and orange circles represent candidates identified using proximal tubular cells. Numbers indicate genes unique to each analysis or shared between the two analyses. For the AKI-to-CKD transition, 537 candidates were unique to the all-cell analysis, 883 were shared, and 3,573 were unique to the proximal-tubule-restricted analysis. (B) Representative GO Biological Process terms shared between the all-cell and proximal tubule–restricted AKI-to-CKD deletion analyses. Shared biological themes included cell adhesion, localization, signal transduction, regulation of multicellular organismal processes and development, cell migration and locomotion, regulation of responses to stimuli, antigen processing and presentation, and regulation of cell differentiation. In (B), dot color represents -log10 of the FDR-adjusted *P* value, and dot size represents the number of genes contributing to each enriched GO term. Redundant GO terms were consolidated to display representative biological themes.

**Supplementary Figure 2.**
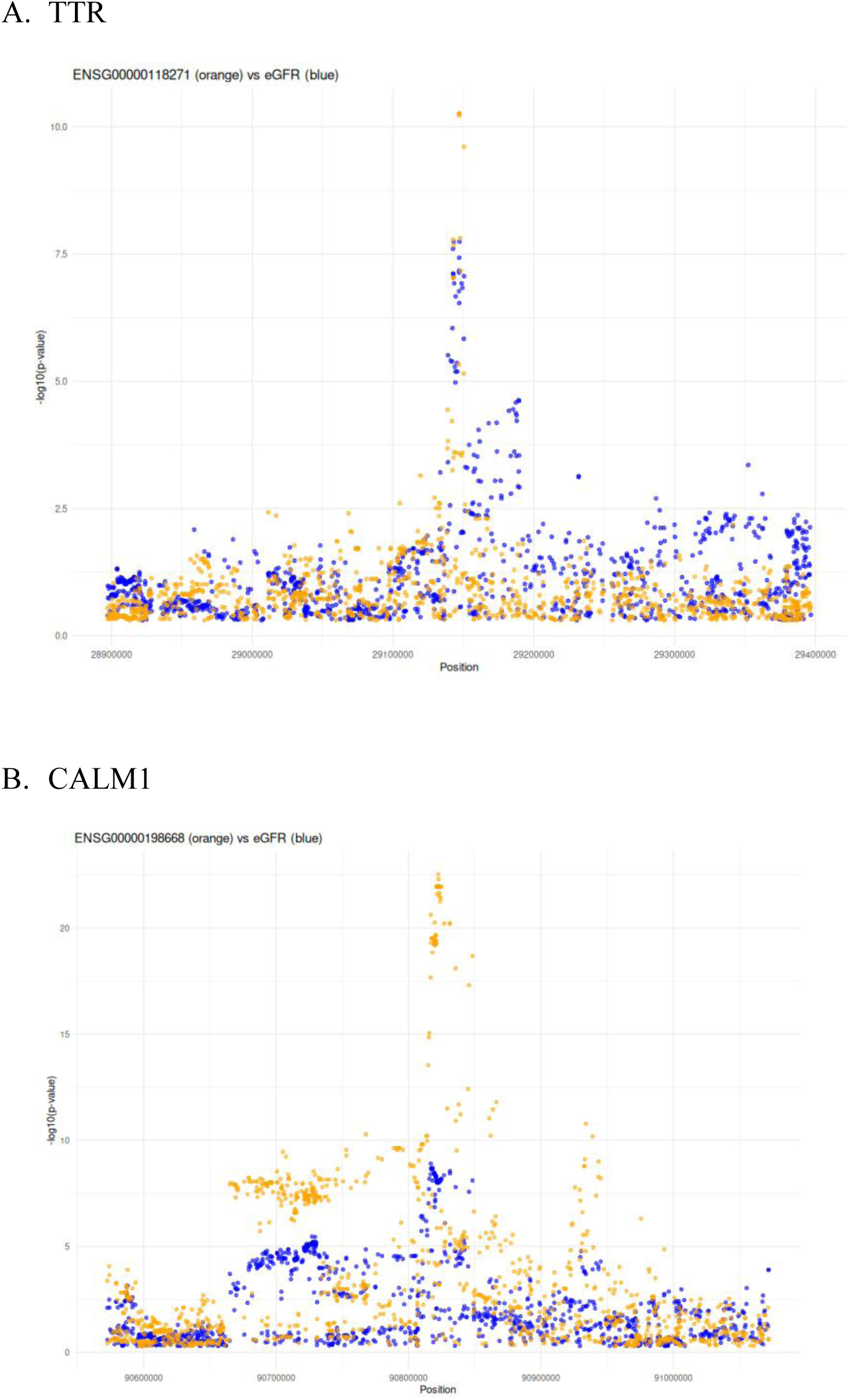
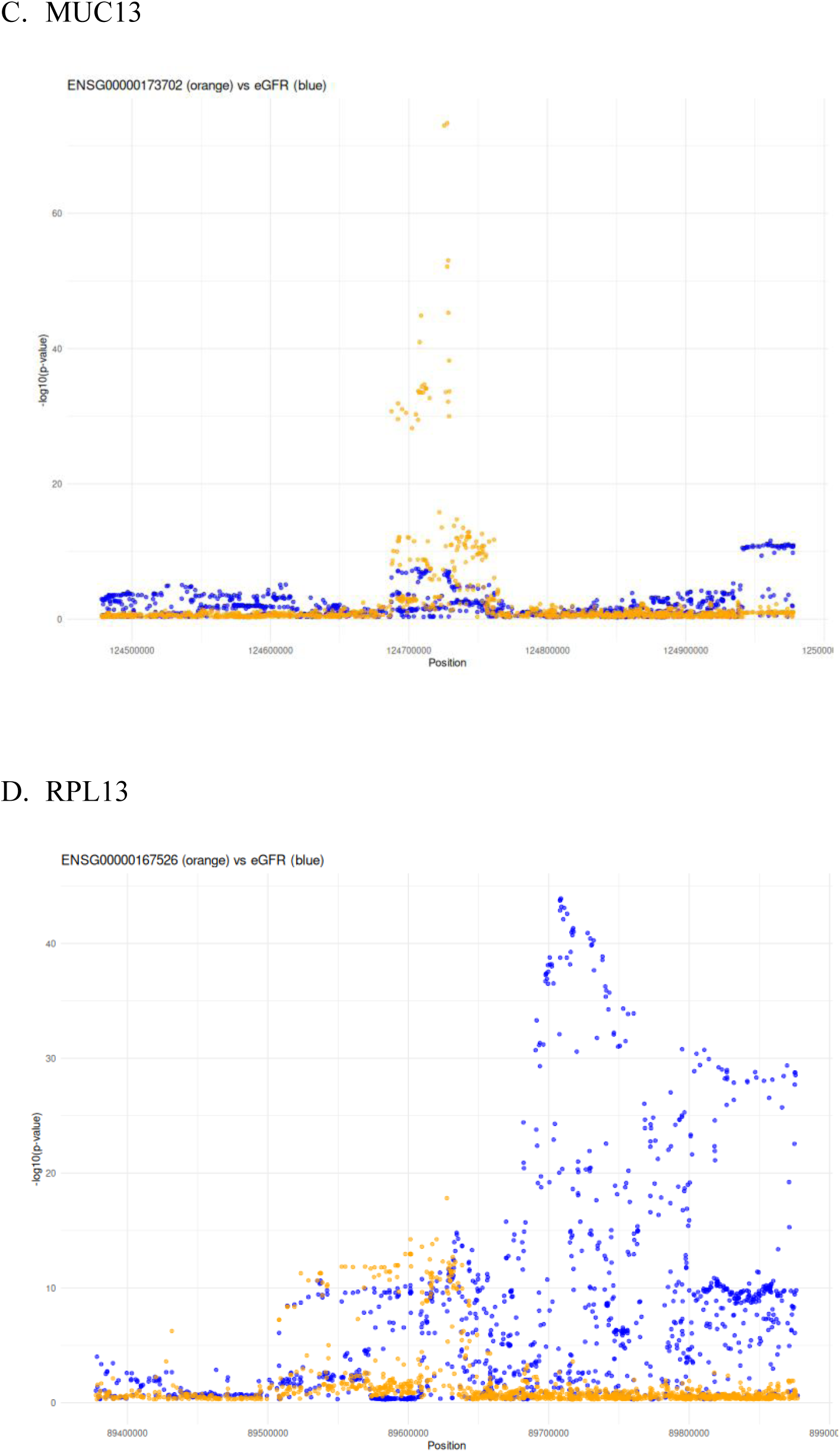
Regional association plots for additional REN-former– prioritized genes evaluated by colocalization analysis. Regional association plots showing tubular expression quantitative trait locus (eQTL) associations and estimated glomerular filtration rate (eGFR) genome-wide association study (GWAS) associations across the genomic regions surrounding (A) *TTR*, (B) *CALM1*, (C) *MUC13*, and (D) *RPL13*. Orange points represent associations with tubular gene expression, and blue points represent associations with eGFR. The overlapping association patterns for *TTR* and *CALM1* were consistent with a shared underlying variant, whereas *MUC13* and *RPL13* showed more limited evidence of colocalization.

**Supplementary Figure 3.**
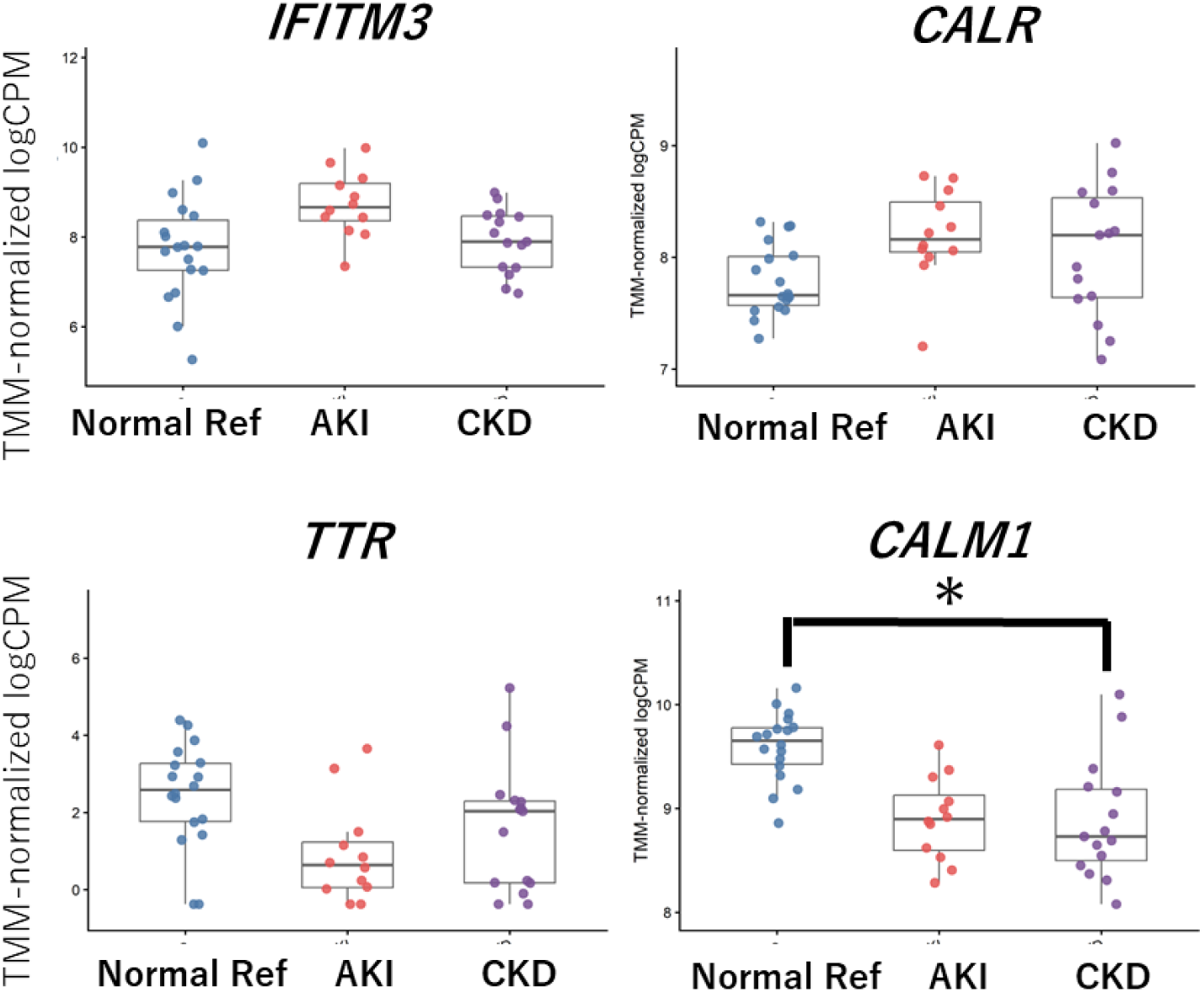
Participant-level proximal-tubule expression of REN-former-prioritized genes supported by colocalization analysis. Participant-level expression of *IFITM3, CALR, TTR,* and *CALM1* in proximal tubular cells from Normal Reference, AKI, and CKD samples in GSE183276. Proximal-tubule UMI counts were aggregated for each participant and converted to TMM-normalized logCPM values. Each point represents one participant; box plots show the median and interquartile range. Differential-expression statistics were obtained from the existing participant-level edgeR analysis. Genes were considered differentially expressed when the false discovery rate was <0.05 and the absolute log2 fold change was ≥0.5. *CALM1* was significantly lower in AKI and CKD than in Normal Reference samples, whereas *IFITM3, CALR,* and *TTR* did not meet the prespecified criteria.

**Supplementary Figure 4.**
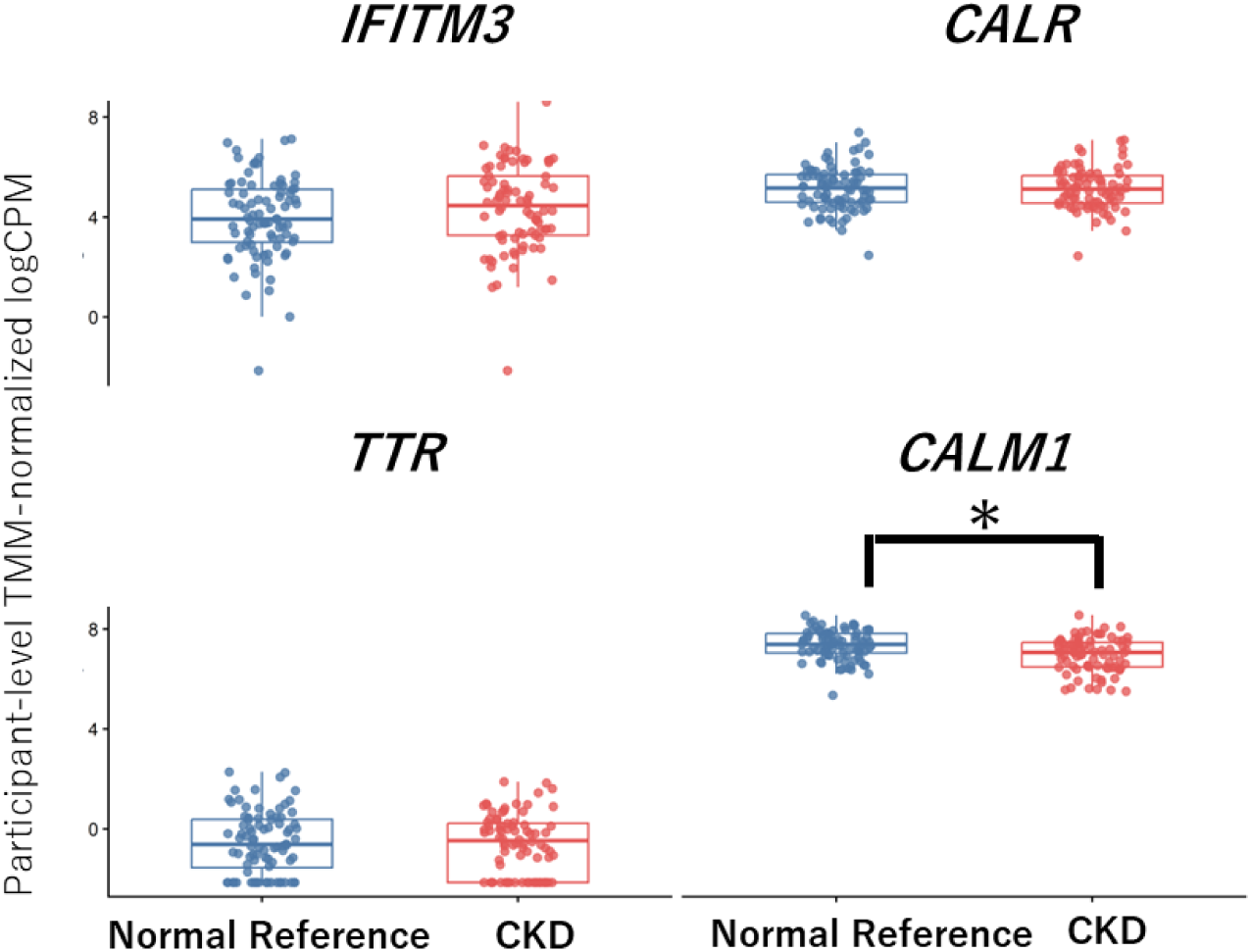
snRNA-seq replication analysis of colocalized candidate genes in non-overlapping KPMP participants. Participant-level expression of *IFITM3, CALR, TTR,* and *CALM1* in broad proximal-tubule cells from 84 Normal Reference and 80 CKD participants. Raw counts were aggregated for each participant and visualized as TMM-normalized logCPM. Each point represents one participant, and boxes indicate the median and interquartile range. *CALM1* was significantly lower in CKD, whereas *IFITM3, CALR,* and *TTR* did not meet the four-gene FDR threshold of 0.05.

