## Supplementary Table Legends for "REN-former prioritizes candidate regulators of kidney disease-state transitions through single-cell foundation modeling and human genetics"

**Supplementary Table 1. Entrez-mapped candidate genes identified by REN-former deletion perturbation analysis for transitions toward AKI- and CKD-like states.**

Complete lists of directionally significant candidate genes that were successfully mapped to Entrez Gene identifiers are shown for (A) the Normal Reference-to-AKI transition, (B) the AKI-to-CKD transition, and (C) the Normal Reference-to-CKD transition. Analyses were restricted to proximal tubular cells. Candidate genes were defined as genes whose deletion induced a positive shift toward the goal state with a false discovery rate–adjusted P value <0.05.

**Supplementary Table 2. Biological processes associated with REN-former deletion perturbation analysis of the AKI-to-CKD transition.**

Significantly enriched nonredundant Gene Ontology Biological Process terms and their associated genes are shown for genes whose in silico deletion induced a positive transcriptomic shift from the AKI-like state toward the CKD-like state. The analysis was restricted to proximal tubular cells. Candidate genes were defined by a positive shift toward the goal state and a false discovery rate–adjusted P value <0.05.

**Supplementary Table 3. Biological processes associated with REN-former overexpression perturbation analyses for transitions toward CKD-like states.**

Significantly enriched nonredundant Gene Ontology Biological Process terms and their associated genes are shown for (A) the Normal Reference-to-CKD transition and (B) the AKI-to-CKD transition. Analyses were restricted to proximal tubular cells. Candidate genes were defined as genes whose overexpression induced a positive shift toward the goal state with a false discovery rate–adjusted P value <0.05.

**Supplementary Table 4. Entrez-mapped candidate genes identified by REN-former overexpression perturbation analysis for transitions toward CKD-like states.**

Complete lists of directionally significant candidate genes that were successfully mapped to Entrez Gene identifiers are shown for (A) the Normal Reference-to-CKD transition and (B) the AKI-to-CKD transition. Analyses were restricted to proximal tubular cells. Candidate genes were defined as genes whose overexpression induced a positive shift toward the goal state with a false discovery rate–adjusted P value <0.05.

**Supplementary Table 5. Biological processes associated with REN-former reverse perturbation analyses from a CKD-like toward a Normal Reference–like state.**

Significantly enriched nonredundant Gene Ontology Biological Process terms and their associated genes are shown for (A) gene deletion and (B) gene overexpression perturbation analyses. Analyses were restricted to proximal tubular cells. Candidate genes were defined as genes whose perturbation induced a positive shift toward the Normal Reference goal state with a false discovery rate–adjusted P value <0.05.

**Supplementary Table 6. Entrez-mapped candidate genes identified by REN-former reverse perturbation analysis for shifts from CKD-like toward Normal Reference–like states.**

Complete lists of directionally significant candidate genes that were successfully mapped to Entrez Gene identifiers are shown for (A) gene deletion and (B) gene overexpression analyses of the CKD-to-Normal Reference transition. Analyses were restricted to proximal tubular cells. Candidate genes were defined as genes whose in silico perturbation induced a positive shift toward the predefined Normal Reference–like goal state with a false discovery rate–adjusted *P* value <0.05.

**Supplementary Table 7. Complete SMR results for REN-former-prioritized gene–analysis pairs.**

Summary-data-based Mendelian randomization (SMR) was performed using tubulointerstitial expression quantitative trait locus data and eGFR genome-wide association study summary statistics after excluding genes within the major histocompatibility complex region. The Bonferroni-corrected significance threshold was $P < 0.05/199$. "HEIDI passed" indicates a HEIDI $P$ value > 0.05. Directional concordance was defined as an SMR effect consistent with the corresponding REN-former-predicted perturbation direction: a positive SMR effect for CKD-to-Normal Reference overexpression or Normal Reference-to-CKD deletion, and a negative SMR effect for CKD-to-Normal Reference deletion or Normal Reference-to-CKD overexpression. Genes meeting the SMR threshold, passing the HEIDI test, and showing directional concordance were selected for colocalization analysis. A gene may appear more than once if it was prioritized in multiple REN-former perturbation analyses. The table contains 221 gene–analysis pairs representing 193 unique genes; candidates without an available SMR result are not represented.
