## Supplementary Methods for "REN-former prioritizes candidate regulators of kidney disease-state transitions through single-cell foundation modeling and human genetics"

**GSE183276 dataset and disease-state definitions**

We used the processed human kidney single-cell RNA-sequencing dataset GSE183276 from the Gene Expression Omnibus. The dataset contained 109,741 cells and 37,080 genes and included participant identifiers, disease groups, kidney cell types, kidney structures, and the original uniform manifold approximation and projection coordinates. The dataset included living-donor kidney biopsy samples from 18 Normal Reference participants contributed by the CZI-HCA and kidney biopsy samples from 12 AKI and 15 CKD participants enrolled in the KPMP. The AKI group included participants with KDIGO stage 1 ($n$ = 1), stage 2 ($n$ = 5), or stage 3 ($n$ = 6) AKI. Histopathological findings were heterogeneous; acute tubular injury was present in 10 of the 12 biopsies, including one with acute interstitial nephritis.

The dataset comprised 21,650 cells from 18 Normal Reference kidneys, 35,777 cells from 12 kidneys with acute kidney injury, 43,076 cells from kidneys with diabetic kidney disease, and 9,238 cells from kidneys with hypertensive chronic kidney disease. Diabetic kidney disease and hypertensive chronic kidney disease were combined into a single CKD group, resulting in 21,650 Normal Reference, 35,777 AKI, and 52,314 CKD cells. Disease groups were defined using the original condition.long metadata field.

We used the processed expression data, kidney-structure labels, cell-type annotations, and UMAP coordinates provided by the original study and did not reprocess the raw sequencing reads or perform new cell-type annotation. Cells labeled as proximal tubules were used for proximal-tubule-specific analyses, whereas analyses of all kidney cells were performed without filtering by kidney structure.

**Participant-level data partitioning for REN-former**

The dataset was partitioned at the participant level into mutually exclusive training ($n$ = 34; 14 Normal Reference, 9 AKI, and 11 CKD), validation ($n$ = 5; 2 Normal Reference, 2 AKI, and 1 CKD), and test ($n$ = 6; 2 Normal Reference, 1 AKI, and 3 CKD) sets. Participant assignments were explicitly specified using the patient metadata field, and all cells from each participant were retained within a single partition. No participant identifiers overlapped among the three sets. The test set was used for final model evaluation and in silico perturbation analyses.

**Detailed participant-level differential-expression and Hallmark pathway analyses**

Integer UMI counts were summed across proximal-tubule cells for each participant to generate pseudobulk profiles, with each participant treated as one biological sample. Participants with fewer than 20 proximal-tubule cells were excluded. The analysis included 24,985 cells from 45 participants: 18 Normal Reference, 12 AKI, and 15 CKD. Diabetic kidney disease and hypertensive CKD were combined into the CKD group.

Differential expression was analyzed using edgeR.[23, 24] Genes were filtered using filterByExpr, with a minimum count of 10 and a minimum total count of 15, and library sizes were normalized using the trimmed mean of M-values method. Robust quasi-likelihood negative-binomial models included disease group and, where possible, sex. Data source was not included because of its close association with disease group. Pairwise comparisons were performed between Normal Reference and AKI, Normal Reference and CKD, and AKI and CKD. $P$ values were adjusted using the Benjamini–

Hochberg method, and differential expression was defined as FDR <0.05 and absolute log2 fold change ≥0.5.

For visualization and pathway analysis, positive log2 fold changes indicated higher expression in Normal Reference for the Normal Reference-versus-AKI and Normal Reference-versus-CKD comparisons and higher expression in AKI for the AKI-versus-CKD comparison. Reversing the contrast direction changed only the sign of the fold change and did not affect the *P* or FDR values.

Preranked Hallmark gene set enrichment analysis was performed using fgseaMultilevel in fgsea version 1.38.0. Human Hallmark gene sets were obtained using msigdbr version 26.1.0 (species = "Homo sapiens", collection = "H").[25, 26] Genes were ranked using the direction of expression change and the unadjusted *P* value from edgeR. Genes with missing values were excluded, and duplicate gene symbols were removed. GSEA was performed using gene symbols, gene-set sizes of 15–500, eps = 0, scoreType = "std", and a random seed of 183276. Hallmark gene sets with Benjamini–Hochberg-adjusted *P* <0.05 were considered significant.

**Detailed Gene Ontology enrichment analysis**

GO Biological Process enrichment analysis was performed separately for each REN-former transition, perturbation type, and cell population.[27] Candidate genes were required to show both a positive predicted shift toward the prespecified goal state (Shift_to_goal_end > 0) and an FDR below 0.05 (Goal_end_FDR < 0.05). Genes with missing or nonfinite values were excluded. All genes meeting these criteria were included without restricting the analysis to a fixed number of top-ranked genes.

The background for each enrichment analysis comprised all unique genes statistically evaluated in the corresponding perturbation analysis. Duplicate gene symbols were resolved by retaining the result with the lowest FDR, followed by the largest shift value and, if still tied, the earliest worksheet row. Gene symbols were converted to Entrez Gene identifiers using org.Hs.eg.db version 3.23.1. When a gene symbol mapped to multiple Entrez identifiers, all mappings were recorded and the lowest numerical identifier was used. Unmapped genes were excluded from enrichment analysis but retained in the mapping audit. All mapped candidate genes were confirmed to be present in the corresponding background set.

GO Biological Process enrichment was tested using clusterProfiler::enrichGO version 4.20.0 with a hypergeometric over-representation test (ont = "BP", keyType = "ENTREZID", minGSSize = 3, and maxGSSize = 5000).[28] Initial *P*- and *q*-value cutoffs were set to 1 to retain the complete results.

Redundant significant terms were grouped using the clusterProfiler simplify function with Wang semantic similarity and a cutoff of 0.7.[29] Within each group, the term with the lowest adjusted *P* value was retained as the representative term. Up to 10 representative terms were displayed, ordered by adjusted *P* value and then by the number of overlapping genes. No terms were manually substituted.

For comparisons between all kidney cells and proximal tubular cells, enrichment analysis was performed separately for each cell population using its corresponding background. A GO term was considered shared when the same GO identifier was significant in both analyses. Gene overlap was summarized using the number of shared genes and the Jaccard index.

**Additional details of the non-overlapping KPMP expression analysis**

Broad proximal tubules were defined using Cell Ontology identifiers CL:0002306, CL:4030009, CL:4030010, and CL:4030011. Integer counts from adata.raw.X were summed by participant to generate pseudobulk profiles. Genes were filtered using filterByExpr, and library sizes were normalized using the TMM method. Differential expression was analyzed using robust dispersion estimation and robust quasi-likelihood generalized linear models in edgeR. Log2 fold changes were defined as CKD relative to Normal Reference, such that positive values indicated higher expression in CKD; this was opposite to the Normal Reference-referenced convention used for the GSE183276 analysis.

**Software and statistical analysis**

Single-cell data processing, REN-former training and evaluation, and in silico perturbation analyses were performed in Python using Geneformer and related libraries. Statistical analyses were performed in R version 4.6.0. Participant-level differential-expression analysis used edgeR; Hallmark gene set enrichment analysis used fgsea version 1.38.0 and msigdbr version 26.1.0; and GO analysis used clusterProfiler version 4.20.0 and org.Hs.eg.db version 3.23.1. SMR and colocalization analyses used SMR version 1.3.2 and coloc version 5.2.3, respectively.

Unless otherwise specified, *P* values were adjusted using the Benjamini–Hochberg method, and an adjusted *P* value below 0.05 was considered significant. The Bonferroni-corrected significance threshold for the SMR analysis was 0.05/199.
